# Molecular evolution of the ependymin-related gene *epdl2* in African weakly electric fish

**DOI:** 10.1101/2022.07.13.499928

**Authors:** Mauricio Losilla, Jason R. Gallant

## Abstract

Gene duplication and subsequent molecular evolution can give rise to taxon-specific gene specializations. In a previous study, we found evidence that African weakly electric fish (Mormyridae) may have as many as three copies of the *epdl2* gene, and the expression of two *epdl2* genes is correlated with electric signal divergence. *Epdl2* belongs to the ependymin-related family (EPDR), a functionally diverse family of secretory glycoproteins. In this study, we first describe vertebrate EPDR evolution and then present a detailed evolutionary history of *epdl2* in Mormyridae with emphasis on the speciose genus *Paramormyrops*. Using Sanger sequencing, we confirm three apparently functional *epdl2* genes in *P. kingsleyae*. Next, we developed a nanopore-based amplicon sequencing strategy and bioinformatics pipeline to obtain and classify full-length *epdl2* gene sequences (N = 34) across Mormyridae. Our phylogenetic analysis proposes three or four *epdl2* paralogs dating from early *Paramormyrops* evolution. Finally, we conducted selection tests which detected positive selection around the duplication events and identified ten sites likely targeted by selection in the resulting paralogs. These sites’ locations in our modeled 3D protein structure involve four sites in ligand binding and six sites in homodimer formation. Together, these findings strongly imply that *epdl2* genes display signatures of selection-driven functional specialization after tandem duplications in the rapidly speciating *Paramormyrops*. Considering previous evidence, we propose that *epdl2* may contribute to electric signal diversification in mormyrids, an important aspect of species recognition during mating.

## Introduction

Gene duplication is an important source of raw genetic material for evolution to act on (Ohno 1970). While gene duplications occur frequently (Anderson and Roth 1977; Bailey et al. 2002; Lynch et al. 2008; Lipinski et al. 2011), the fate of many gene duplicates is pseudogenization (Lynch and Conery 2000). In some cases, gene duplicates take on new or specialized functions through molecular evolution (Taylor and Raes 2004; Conant and Wolfe 2008; Chapal et al. 2019), contributing to the emergence of species-specific characteristics and taxon diversification (Zhang 2003; Magadum et al. 2013). Weakly electric fishes provide elegant examples of this phenomenon: duplicate sodium channel genes resulting from the teleost-specific whole genome duplication were independently neofunctionalized in electric organs convergently in Neotropical gymnotiformes and African mormyroids (Zakon et al. 2006; Arnegard, Zwickl, et al. 2010). Within mormyroids, duplicate potassium channel genes resulting from the teleost-specific whole genome duplication, exhibit a similar pattern of neofunctionalization in electric organs (Swapna et al. 2018). By leveraging the well-characterized structure-function relationship of ion channels, these studies have illustrated how neofunctionalization of gene duplicates can be a powerful evolutionary phenomenon.

African weakly electric fish (Mormyridae) are among the most rapidly speciating clades of ray-finned fishes (Carlson et al. 2011; Rabosky et al. 2013). Often, mormyrid species are most reliably discriminated by the electric organ discharges (EODs) they generate (Hopkins 1981; Bass 1986; Alves-Gomes and Hopkins 1997; Sullivan et al. 2000; Arnegard et al. 2005). This is well illustrated by the genus *Paramormyrops*, whose rapid and recent diversification in West-Central Africa has given rise to more than 20 species (Sullivan et al. 2002; Sullivan et al. 2004; Lavoué et al. 2008; Peterson et al. 2022). Within *Paramormyrops*, EOD waveform evolution greatly outpaces that of morphology, size, and trophic ecology (Arnegard, McIntyre, et al. 2010). Mormyrid EODs are a central component of their electrolocation (Lissmann and Machin 1958; von der Emde et al. 2008) and communication (Möhres 1957; Kramer 1974) behaviors, and principally vary in the number of EOD phases present and their duration. In a recent study, we identified gene expression correlates for variable EOD features among *Paramormyrops* species (Losilla et al. 2020), and demonstrated that two copies of the gene *ependymin-like* 2 (*epdl2*) were the most highly differentially expressed genes between biphasic and triphasic *Paramormyrops*. In addition, we found reduced *epdl2* expression in individuals with longer EODs in the mormyrid *Brienomyrus brachyistius* (Losilla M, Gallant JR, unpublished data).

*Epdl2* is a member of the ependymin-related family (EPDR), a widespread but understudied family of secretory, calcium-binding glycoproteins. Generally, EPDR proteins change their structural conformation according to calcium concentration (Ganss and Hoffmann 2009), and participate in cell-cell and cell-extracellular matrix interactions (Schwarz et al. 1993; Hoffmann 1994; Pradel et al. 1999). The EPDR family has undergone extensive independent tandem duplications and divergence within metazoans (McDougall et al. 2018), such that EPDR paralogs have been proposed as suitable targets to experimentally test gene subfunctionalization (Suárez-Castillo and García-Arrarás 2007). EPDRs participate in an extraordinarily diverse range of functions, including memory formation and neuroplasticity in fish (Shashoua 1985; Schmidt et al. 1995; Pradel et al. 1999), shell patterning and pigmentation in gastropods (Jackson et al. 2006), intestinal regeneration in sea cucumbers (Zheng et al. 2006), collagen contractibility of human fibroblasts (Staats et al. 2016), and conspecific communication in crown-of-thorns starfish (Hall et al. 2017). We noted in an initial study that the *Paramormyrops kingsleyae* genome contains as many as three *epdl2* genes, whereas the osteoglossiform *Scleropages formosus* has only one, strongly suggesting that *epdl2* underwent duplications during mormyrid evolution. This discovery presents a unique opportunity to study the molecular evolution of an enigmatic protein, expressed in the mormyrid electric organ, that likely underwent linage-specific duplications.

In this study, we begin with an examination of vertebrate EPDR evolution, including *epdl2*. Next, we focus on a detailed molecular evolutionary analysis of *epdl2* in African weakly electric fish, demonstrating that a lineage-specific tandem duplication of *epdl2* genes occurred within mormyrids, specifically near the origin of the *Paramormyrops* genus. Next, we test for signatures of selection on *epdl2* paralogs within mormyrids and among the *Paramormyrops*, identifying ten sites of interest that have experienced positive selection and have evolved at higher ω rates in the lineages with *epdl2* duplications. Finally, we model the 3D structure of a representative Epdl2 protein and propose how the ten sites of interest may affect protein function. This work demonstrates that *epdl2* genes underwent tandem duplications near the origin of the rapidly speciating *Paramormyrops*, primarily distinguishable by their EOD signal (Sullivan et al. 2002; Arnegard and Hopkins 2003; Picq et al. 2020). The resulting *epdl2* paralogs have experienced rapid sequence evolution and specialization. Given our previous work demonstrating strong differential expression of *epdl2* paralogs in electric organs with different signal types, we hypothesize that *epdl2* evolution may play an important role in the evolution of EOD signals.

## Methods

### Evolutionary relationships between EPDR vertebrate genes

We used Genomicus (Nguyen et al. 2022) v03.01 to visualize homology relationships between vertebrate EPDR genes. This version is based on a new, comparative atlas of teleost genomes (Parey et al. 2022), which provided additional non-teleost *epdl* sequences, including *Amia calva* (bowfin). First, we obtained a gene tree for the EPDR family in chordates in Genomicus PhyloView with the EPDR family as reference (Fam016630) and the root species at Chordata. Second, we explored synteny relationships between *epdl* genes to further understand their evolutionary relationships. We used Genomicus PhyloView with bowfin *epdl* as reference to visualize the genomic neighborhood of the *epdl* teleost homologs.

Next, we performed a phylogenetic analysis of selected *epdl* genes proposed by the Genomicus EPDR tree, detailed in Table S1. These included i) Epdl amino acid sequences from all non-teleost vertebrates included in Genomicus v03.01, plus a sequence from *Protopterus annectens* (lungfish) identified via BLAST search, ii) representative amino acid sequences from basal teleost taxa (Osteoglossiformes, Otocephala, and Euteleosteomorpha) for each of the *epdl* homologs identified by the gene tree, and iii) the taxon-specific genes in *Clupea harengus* and in *Paramormyrops kingsleyae* as proposed by the gene tree, plus a related sequence from *B. brachyistius*, identified by BLAST search against its genome. We manually curated several of the gene annotations that supported these amino acid sequences (Table S1 and Additional file 1). We aligned the selected and curated sequences in Geneious 9.1.8 (Biomatters) using Muscle (Edgar 2004) (default settings) and manually inspected the resulting alignment. This alignment had 238 sites, but we trimmed the highly variable N (29 sites) and C (32 sites) termini. The final alignment (177 sites) consisted of the amino acids between (and including) the first and last highly conserved cysteine residues among *epdl* genes. We used this alignment to infer a vertebrate *epdl* gene tree based on Bayesian inference with MrBayes 3.2.7 (Huelsenbeck and Ronquist 2001) through the online portal of NGPhylogeny.fr (Lemoine et al. 2019) (https://ngphylogeny.fr/) (run length: 1 000 000 generations, rates = invgamma, all other options set to default values).

### *epdl2* sequences from Osteoglossiformes genomes

We obtained *epdl2* gene sequences from genome assemblies of *S. formosus, Gymnarchus niloticus* and *B. brachyistius* (Table S2). We identified the *epdl2* genes with BLAST searches against each genome, using *epdl2* genes (blastn) and proteins (tblastn) as queries. For *B. brachyistius*, the *epdl2* queries were the incomplete *epdl2* gene models from the *P. kingsleyae* Nanopore assembly (see below); and once identified, the *B. brachyistius epdl2* gene served as query for the *G. niloticus* and *S. formosus* searches.

### Confirm *epdl2* triplication in *Paramormyrops kingsleyae*

#### *P. kingsleyae* Nanopore assembly

We generated a new *P. kingsleyae* genome assembly based on a single *P. kingsleyae* adult female (Table S3, sample name MSU-160) captured from Apassa Creek, Gabon, West-Central Africa. We refer to this individual as *P. kingsleyae* (APA) throughout this paper. We verified its EOD as triphasic and confirmed its sex by postmortem gonad inspection. We isolated High molecular weight genomic DNA from fin clip tissue using a NanoBind Tissue Big DNA Kit (Circulomics, cat. NB-900-701-01) following the TissueRuptor Homogenizer Protocol (Qiagen, v0.17). Extracted gDNA was verified for concentration and quality using a Nanodrop and Qubit (Thermo Fisher Scientific), and high molecular weight DNA was confirmed on a 0.5% agarose gel. Sequencing libraries were constructed using a Voltrax V2 device (Oxford Nanopore Technologies, ONT) with the Voltrax Sequencing Kit (ONT, VSK-002) following manufacturer’s instructions. Based on previous assembly size of 885 Mb of the *P. kingsleyae* genome (Gallant et al. 2017), we targeted 20x coverage using two R9.4.1 flow cells (ONT). We obtained 2,250,969 reads. Raw reads were corrected for errors and assembled into contigs using Canu (Koren et al. 2017) with default settings. The resulting assembly was 862 Mb with a contig N50 of 446,819 bp. Raw sequencing reads supporting this assembly have been deposited into the SRA with accession number SRR19573669. This genome assembly has been deposited at GenBank under the accession JAMQYA000000000. We refer to this assembly as the Nanopore assembly.

#### *epdl2* paralogs in the *P. kingsleyae* Nanopore assembly

The NCBI-annotated (Release 100) *P. kingsleyae* genome (Gallant et al. 2017) contains three, incomplete *epdl2* genes. We used these sequences, along with flanking genes *otu1b* and *nrg2*, to find the *epdl2* genes in the *P. kingsleyae* Nanopore assembly. All queries mapped to one gene region, where the NCBI annotation suffered from insufficient scaffolding, and the Nanopore assembly contained frame shifts caused by small indels in coding homopolymers. These structural problems impeded complete and accurate *epdl2* annotations in each assembly. As such, we decided to amplify and Sanger-sequence each potential paralog to confirm the presence of three *epdl2* paralogs and to deduce each paralog’s gene structure and coding sequence.

#### PCR amplification and Sanger sequencing of *epdl2* paralogs in *P. kingsleyae*

We designed primers to amplify each *P. kingsleyae epdl2* paralog (Table S4). Their binding sites are 500-1500 bp up or downstream of the start/stop codon of each paralog, where primer binding was predicted to amplify a unique PCR product. For PCR amplifications we used Q5 High-Fidelity DNA Polymerase (NEB, cat. M0491S) and high molecular weight DNA from the same individual used to sequence the Nanopore assembly. We carried out PCRs in 50 μl reactions, with the reagents and conditions detailed in Tables S5 & S6. PCR products were cleaned (Monarch PCR & DNA Cleanup Kit, NEB cat. T1030S) and Sanger-sequenced. For the latter, in addition to the PCR primers, we designed sequencing primers (Table S4) that bind to regions conserved across all three putative paralogs and were spaced by < ~700 bp. Thus, each amplified *epdl2* paralog was sequenced with six or seven Sanger-sequencing reactions. Finally, we used the obtained paralog-specific sequences and the *epdl2* annotations from *B. brachyistius* to produce gene models for each *epdl2* paralog in *P. kingsleyae*.

### *epdl2* sequences across Mormyridae

#### Primer design

We designed primers to target all *epdl2* paralogs in a given mormyrid species (Table S4). As primer-binding regions, we chose the 5’and 3’ UTRs, close to the start and stop codons. Primer design was guided by the Sanger-sequenced *P. kingsleyae epdl2* paralogs, our *B. brachyistius epdl2* sequence, and whole genome sequencing scaffolds from species across Mormyridae (Table S7). We assembled the target UTRs from whole genome sequencing scaffolds i) returned from BLAST searches (tblastn) of *P. kingsleyae* and *B. brachyistius* Epdl2 proteins against the scaffolds, or ii) that mapped to the *P. kingsleyae* or *B. brachyistius epdl2* genomic region.

#### Sample collection, DNA extraction, PCR amplification and *epdl2* sequencing

Individual mormyrid specimens (Table S3) were euthanized with an overdose of MS-222, and one or more paired fins from each were clipped and preserved in 95% ethanol. Field captured specimens were caught using previously described methods (Gallant et al. 2011; Picq et al. 2020). The remaining specimens were obtained from the pet trade, or generously provided by other laboratories. All methods described conform to protocols approved by Michigan State University Institutional Animal Use and Care Committee (IACUC).

We extracted DNA from the ethanol-preserved fin clips, from one individual of each of our studied species, with the DNeasy Blood & Tissue Kit (Qiagen, cat. 69504). Then we targeted all *epdl2* paralogs for PCR amplification in 25 μl reactions, with primer- and species-dependent reagents (Table S8) and conditions (Table S9). The resulting amplicons were multiplexed and sequenced on an ONT MinION device with a R9.4.1 flow cell. The multiplexed sequencing library was prepared with ONT kits: Native Barcoding Expansion 1-12 (cat. EXP-NBD104) & 13-24 (cat. EXP-NBD114), and Ligation Sequencing Kit (cat. SQK-LSK109), following manufacturer’s directions.

### Analysis of Nanopore reads

#### Read processing

Reads were base-called and quality-filtered (Q > 7) with the high accuracy model from Guppy 4.2.3+f90bd04 (ONT), and barcoded reads passing filtering were demultiplexed with Guppy 4.4.1+1c81d62. We uploaded these reads to the NCBI SRA (Table S3) and performed all subsequent steps on these demultiplexed reads.

We modified the amplicon subcommand of seqkit 0.15.1 (Shen et al. 2016) to identify all amplicons in each read. Next, using a custom pipeline consisting of seqkit, Nanofilt 2.71 (De Coster et al. 2018) and custom code, we extracted the smallest amplicon greater than 1000 bp. Extracted amplicons were then subjected to one additional round of size (1200-2400 bp) and quality (Q > 14) filtering with NanoFilt. We generated quality control summaries and plots with Nanoplot 1.33.1 (De Coster et al. 2018).

#### Identification of *epdl2* genes

The following bioinformatic procedure is summarized in Fig. 1. The filtered amplicons from each sample were clustered using cd-hit 4.8.1 (Li and Godzik 2006). A preliminary analysis of the data revealed a key parameter is cd-hit’s sequence identity threshold (c, range 0-1). We noticed in *P. kingsleyae*, where the number of *epdl2* copies were known through Sanger sequencing, that relatively low values of c lumped amplicons from different genes into the same cluster (underclustering), whereas relatively high values split amplicons from the same gene into different clusters (overclustering). To mitigate this issue, for each sample, we ran cd-hit with c values from 0.84 to 0.91 in 0.01 increments and selected the c value that produced the best clustering. We chose this range because the selected c value from every sample was higher than 0.84 and lower than 0.91. The c value that generated the best clustering was chosen as follows: we selected the c value that produced the largest number of supported clusters (we considered a cluster to be supported if it contained >10% of the amplicons). If more than one c value met this criterion, we chose the c value that classified the largest number of amplicons into supported clusters. If more than one c value met this, we selected the largest c value. Once a c value was selected, we generated consensus sequences for each cluster and manually inspected them for overclustering, which in our dataset manifested as consensus sequences from different clusters with greater than 99.45% sequence identity. In these few cases, we reapplied the selection criteria to choose a new c value that produced fewer supported clusters than the discarded c value.

**Fig. 1.**
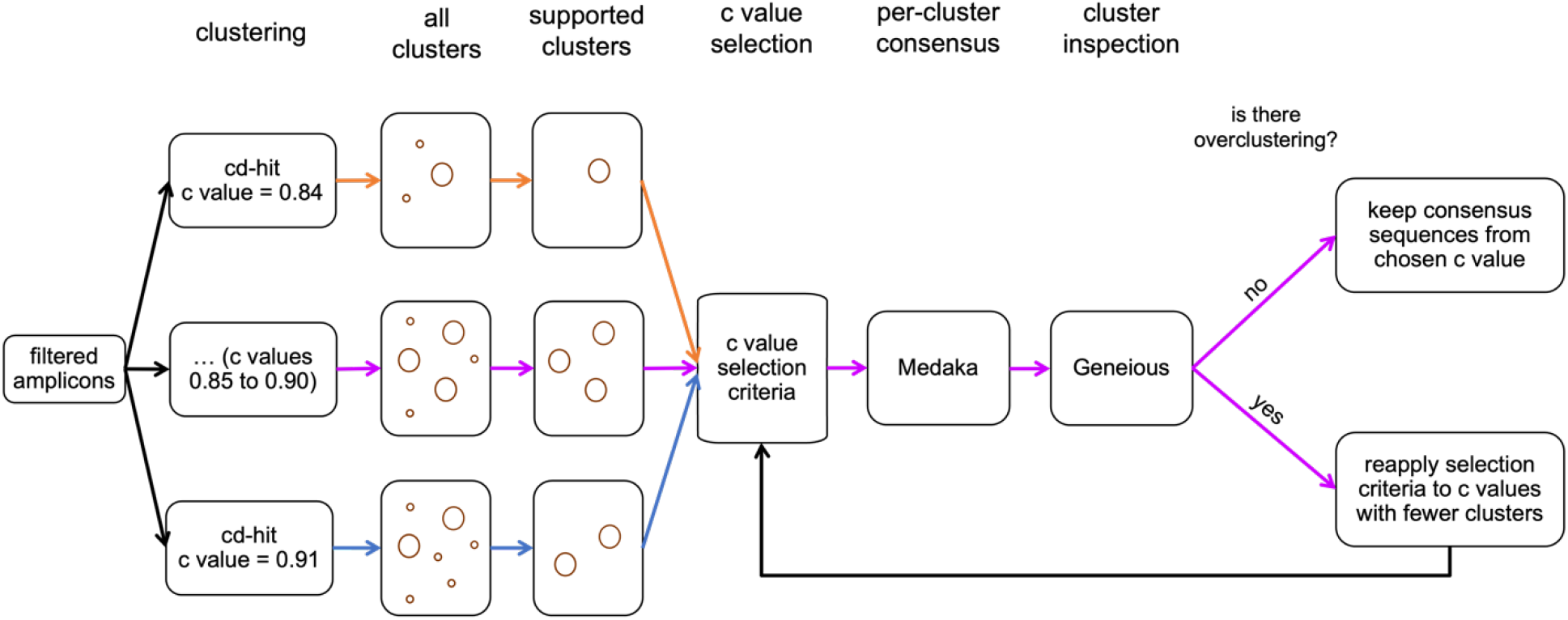
Graphical summary of the bioinformatic pipeline we leveraged to identify *epdl2* genes in the filtered amplicons from each species, see Methods for details. Same-colored connector arrows represent an analysis with a specific value for c (cd-hit’s sequence identity threshold parameter, range analyzed 0.84-0.91). Magenta connector arrows represent the analysis with the c value chosen with the selection criteria.

Following clustering with cd-hit, we obtained consensus sequences from each supported cluster with Medaka 1.2.1 (smolecule module) (https://github.com/nanoporetech/medaka), and manually inspected the consensus sequences in Geneious by aligning amplicons to the *epdl2* sequences of *B. brachyistius* and *P. kingsleyae*. This allowed us to identify and remove nonspecific amplicons. Finally, for each aligned consensus sequence, we manually annotated CDSs and edited homopolymer sequences in CDS4 to keep the sequence in frame. The source code for this pipeline can be found at https://github.com/msuefishlab/molec_evol_epdl2

#### *epdl2* gene tree and *epdl2* duplication history

We built an *epdl2* gene tree to infer the *epdl2* duplication history in Mormyridae. We aligned all the mormyrid *epdl2* CDSs and introns in Geneious using Muscle (default settings) followed by manual inspection. We excluded the *epdl2* genes of *Gymnarchus niloticus* and *Scleropages formosus* because their very long introns 1 and 2 could not be aligned reliably. We used this alignment to infer a mormyrid *epdl2* gene tree based on Bayesian inference, following the same procedure employed in the inference of the vertebrate *epdl* gene tree. In addition, we inferred a mormyrid *epdl2* gene tree with the maximum likelihood (ML) criterion to ensure the tree’s robustness. For the ML tree, we used the online portal of PhyML 3.3 (Guindon et al. 2010) (http://www.atgc-montpellier.fr/phyml). The best-fitting nucleotide substitution model was determined by Smart Model Selection (SMS) 1.8.4 (Lefort et al. 2017) using the Akaike Information Criterion (AIC), and we estimated branch support with 10000 bootstrap replicates. The substitution model K80 + G best fitted the alignment of the *epdl2* coding regions, thus the *epdl2* gene tree was inferred with the model K80 + G + F.

### Selection Tests

In order to test for signatures of natural selection, we extracted CDSs from all the sequences obtained from Nanopore sequencing, as well as additional osteoglossiform sequences (Table S2). Homologous codons were aligned by backtranslating aligned protein sequences on the online portal of TranslatorX (Abascal et al. 2010) (http://translatorx.co.uk/) using the Prank algorithm (Löytynoja and Goldman 2005). Then we manually refined alignment gaps and removed stop codons.

Some selection tests use a user-supplied gene tree topology. For this, we used the topology of our mormyrid *epdl2* gene tree with the addition of *S. formosus* and *G. niloticus* as basal taxa, as supported by topologies derived from CDSs and from amino acid sequences (not shown), and well-established taxonomic relationships within Osteoglossiformes. We call this the osteoglossiform *epdl2* gene tree (Fig. 6).

We ran five tests, using HyPhy (Kosakovsky Pond et al. 2005; Kosakovsky Pond et al. 2020) via their online portal (Weaver et al. 2018) (http://www.datamonkey.org), except where indicated. First, we looked for evidence of gene conversion in the alignment of osteoglossiform *epdl2* CDSs with the method GARD 0.2 (Kosakovsky Pond et al. 2006). We ran this analysis three times, each with one of the three options available for site-to-site variation (none, general discrete, beta-gamma), and in every run the run mode was set to normal and rate classes was set to 4 (default).

We then used a local installation of HyPhy 2.5.30 to run RELAX 4.0 (Wertheim et al. 2015) (synonymous rate variation allowed from site to site, and omega rate classes = 2), to test whether the strength of selection on *epdl2* changed (relaxed or intensified) in the branches with *epdl2* duplications relative to the mormyrid branches without *epdl2* duplications (Fig. 6). After this, we employed the branch-test aBSREL 2.2 (Smith et al. 2015) to identify branches that experienced positive selection in the time between the *epdl2* duplication events and the subsequent divergence into extant *epdl2* paralogs (Fig. 6). We limited this analysis to these branches i) to conserve statistical power, since this test corrects for multiple testing, ii) because the RELAX test suggested that the strength of selection has intensified in the duplicated lineages.

Finally, we attempted to identify sites (codons) of interest in *epdl2* genes. First, we used the site-test MEME 2.1.2 (Murrell et al. 2012) to detect which sites in *epdl2* have been subjected to positive selection. Second, we used Contrast-FEL 0.5 (Kosakovsky Pond et al. 2021) to investigate which sites in *epdl2* may be evolving at different rates between the mormyrid lineages with vs without *epdl2* duplications (Fig. 6). The sites we are most interested in are those identified by both methods. All sites we report are based on the numbering produced by these tests (which in turn are based on the homologous codon alignment), and thus they do not represent amino acid positions for any specific Epdl2 protein.

### Structural predictions of Epdl2

We chose the Epdl2 amino acid sequence of *Paramormyrops* sp. SZA as a representative mormyrid Epdl2 protein to explore structural features common in EPDR proteins. We selected this sequence because it is sister to the sequences with duplicated genes in the mormyrid *epdl2* gene tree. We searched for a signal peptide on the online portal of SignalP 5.0 (Almagro Armenteros et al. 2019) (www.cbs.dtu.dk/services/SignalP/), we identified N-Glycosylation sites on the web implementation of NetNGlyc 1.0 (Gupta and Brunak 2002) (www.cbs.dtu.dk/services/NetNGlyc/), and we predicted a 3D structure of the Epdl2 protein with its signal peptide removed using the I-TASSER server (Roy et al. 2010; Yang et al. 2015; Yang and Zhang 2015) (https://zhanglab.dcmb.med.umich.edu/I-TASSER/). Finally, we employed ChimeraX 1.3 (Pettersen et al. 2021) on the resulting PDB model to predict protein secondary structures, annotate protein features, and highlight sites of interest.

## Results

### Evolutionary relationships between EPDR vertebrate genes

The Genomicus EPDR tree (Fig. 2A) proposes that all vertebrate EPDRs evolved from a chordate EPDR whose ortholog has experienced several duplications within Cephalochordates and Tunicates. A duplication in an early Vertebrate ancestor produced two paralogs: *epdr1*, present in all vertebrates (also known as MERP1 in mammals) and *epdl*, which is absent in tetrapods. *Epdl* has undergone tandem duplications in bowfin, and four potential duplications and allegedly five resulting *epdl* paralogs during early teleost diversification. Two of these paralogs were detected each in only one species: *Paramormyrops kingsleyae* (which we tentatively call *epdl3*) and *Clupea harengus* (the existence of which our analysis does not support, see below). The other three genes correspond to the known EPDR genes *epdl1, epdl2*, and *epd*. The last of these is only found in Clupeocephala, and it experienced an additional duplication before the diversification of this clade. Therefore, we refer to the original paralog, before its additional duplication, as *epd*, and call the extant paralogs *epd1* and *epd2* (the first described member of the EPDR family, named *epd*, belongs to the *epd1* paralog). Although we try to minimize nomenclature conflicts, our gene names will differ from some of the computationally assigned names currently found in annotation databases. Opportunities for discrepancies are exacerbated because every one of the widespread *epdl* paralogs has experienced independent duplications in various teleost taxa.

**Fig. 2.**
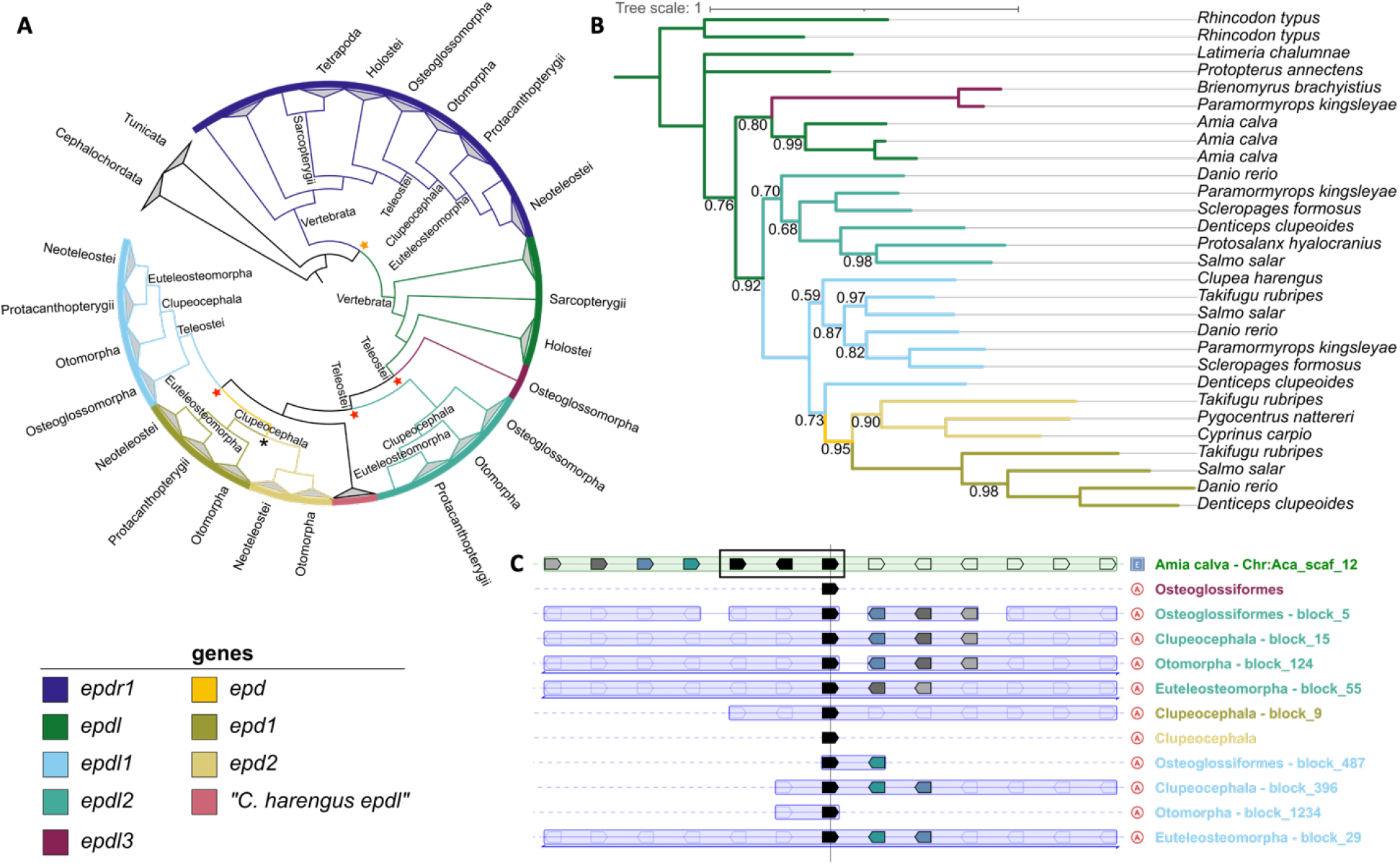
Evolutionary relationships between ependymin-related family (EPDR) vertebrate genes. Gene color legend applies to all panels. A) Chordate EPDR gene tree based on Genomicus v03.01 (Fam016630). Stars represent duplications that led to genes supported by our analysis (orange: ancestral EPDR in early vertebrates, red: *epdl* in early teleosts, black: *epd* in early clupeocephalans). B) Vertebrate *epdl* gene tree based on bayesian inference, only posterior probabilities < 1 are shown. Supported *epdl* genes are color coded in branches. C) Simplified PhyloView alignment (Genomicus v03.01) of the *epdl* teleost homologs in select taxa (rows) aligned to one bowfin *epdl* gene. Black pentagons in each row denote *epdl* genes, including three tandem copies (black rectangle) in bowfin (top row). Pentagons represent the position and orientation of genes syntenic to the *epdl* gene in each taxon. Colored pentagons highlight genes present in a taxon and in the bowfin reference. Taxa labels (right column) are color coded by *epdl* gene.

Our phylogenetic analysis (Table S1, Fig. 2B) clusters most of our selected teleost sequences into three main clades that correspond with each of the three known genes *epdl1, epdl2*, and *epd*, the latter consisting of two distinct clades, *epd1* and *epd2*. Exceptions to this pattern include *Denticeps clupeoides epdl1* with the *epd* genes; the two novel, *epdl3* sequences only known from Osteoglossiformes with the three bowfin *epdl* sequences; and the sequence hypothesized to be a paralog present only in *Clupea harengus* with the *epdl1* sequences.

Synteny analysis reveals that the bowfin *epdl* shares several neighboring genes with both *epdl1* and *epdl2*, but it shares no genes with the remaining teleost *epdl* genes (*epdl3*, *epd1*, and *epd2*) (Fig. 2C). Similar results were observed with the coelacanth *epdl* gene as reference (not shown).

Thus, this analysis greatly clarifies the relationships between the teleost EPDR genes and confirms that the sequences analyzed in the remainder of this study are *epdl2* orthologs.

### *Paramormyrops kingsleyae* has three complete *epdl2* paralogs

We identified a ~40 Kbp region in chr24 in the *P. kingsleyae* Nanopore assembly with three *epdl2* copies in tandem (Fig. 3). Based on this genome sequence, we used PCR and Sanger sequencing to obtain amplicons of the expected sizes (~3200 and ~4000 bp); and confirmed that there are three distinct *epdl2* paralogs in *P. kingsleyae*. Each of our three predicted gene models showcases the six expected CDSs, canonical splicing sites, start and stop codons, and a predicted protein size (213-214 amino acids) of standard EPDR length, with the four conserved cysteine residues, a highly conserved proline residue (McDougall et al. 2018), and high sequence similarity to Epdl2 proteins from other taxa, suggesting that all produce functional proteins. We named these paralogs *epdl2.1, epdl2.2*, and *epdl2.3* (Fig. 3); their respective lengths from start to stop codon are: 1675, 1814, and 1809 bp.

**Fig. 3.**
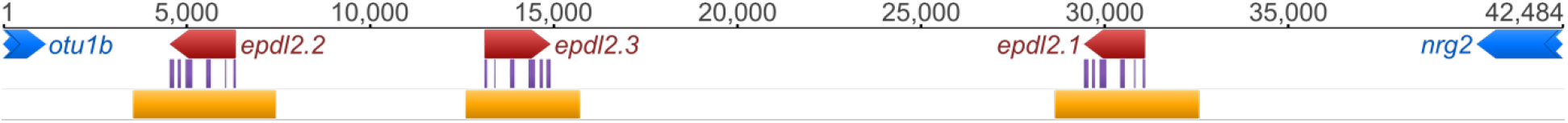
*epdl2* genomic region in *Paramormyrops kingsleyae*, showing three *epdl2* paralogs in tandem. DNA sequence is represented by the black line, numbers above indicate base positions, annotations are shown below the line. Pentagons represent a gene’s position and orientation, flanking genes are depicted in blue, *epdl2* paralogs in red (start to stop codon) and purple (CDSs). Orange blocks mark the Sanger-sequenced regions.

### *epdl2* sequences and copy number across Mormyridae

We developed a Nanopore-based amplicon sequencing strategy to obtain full-length *epdl2* gene sequences across a variety of mormyrid species and used a custom bioinformatics pipeline to identify paralogs. As validation of this approach, we included an additional *P. kingsleyae* sample in our analysis (referred to as *P. kingsleyae* (BAM) after Bambomo Creek, its collection location). Our pipeline found three *epdl2* paralogs in this sample, each shared 99.9% sequence identity from start to stop codon and the same predicted protein sequence to its Sanger sequenced counterpart (*P. kingsleyae* (APA)). When we applied this approach to other mormyrids, we identified a variable number of *epdl2* genes across various mormyrid species, with a median size of 1815 bp (from start to stop codon) (Table S2, which includes GenBank accession numbers). Every *epdl2* gene we identified displays the above mentioned hallmarks of a functional *epdl2* gene. We were unable to obtain *epdl2* sequences in *Paramormyrops* sp. SN2, *Paramormyrops* sp. SN9, *Paramormyrops* sp. TEU, and *Ivindomyrus marchei*. Using this approach, we were able to comprehensively survey *epdl2* sequences among Osteoglossiformes (*Scleropages formosus*), Gymnarchidae (*Gymnarchus niloticus*), and from the main branches of Mormyridae and *Paramormyrops*.

### *Epdl2* duplications occurred early in *Paramormyrops* evolution

Based on these sequences, we generated a mormyrid *epdl2* gene tree. Bayesian inference and ML methods yielded identical trees except for two terminal branches in one of the paralog clades. We present the tree from the former analysis (Fig. 4). Except for a few branches with low support, the topology of this gene tree is congruent with known species relationships outside the *Marcusenius ntemensis* + *Paramormyrops* (Mn+P) clade, where only single-copy *epdl2* genes were detected. This contrasts with the Mn+P clade, where we found all *epdl2* paralogs and a salient disagreement with the species tree: *M. ntemensis* is placed within *Paramormyrops*, although it is well established that these are sister taxa (Sullivan et al. 2002; Lavoué et al. 2003; Sullivan et al. 2004; Peterson et al. 2022).

**Fig. 4.**
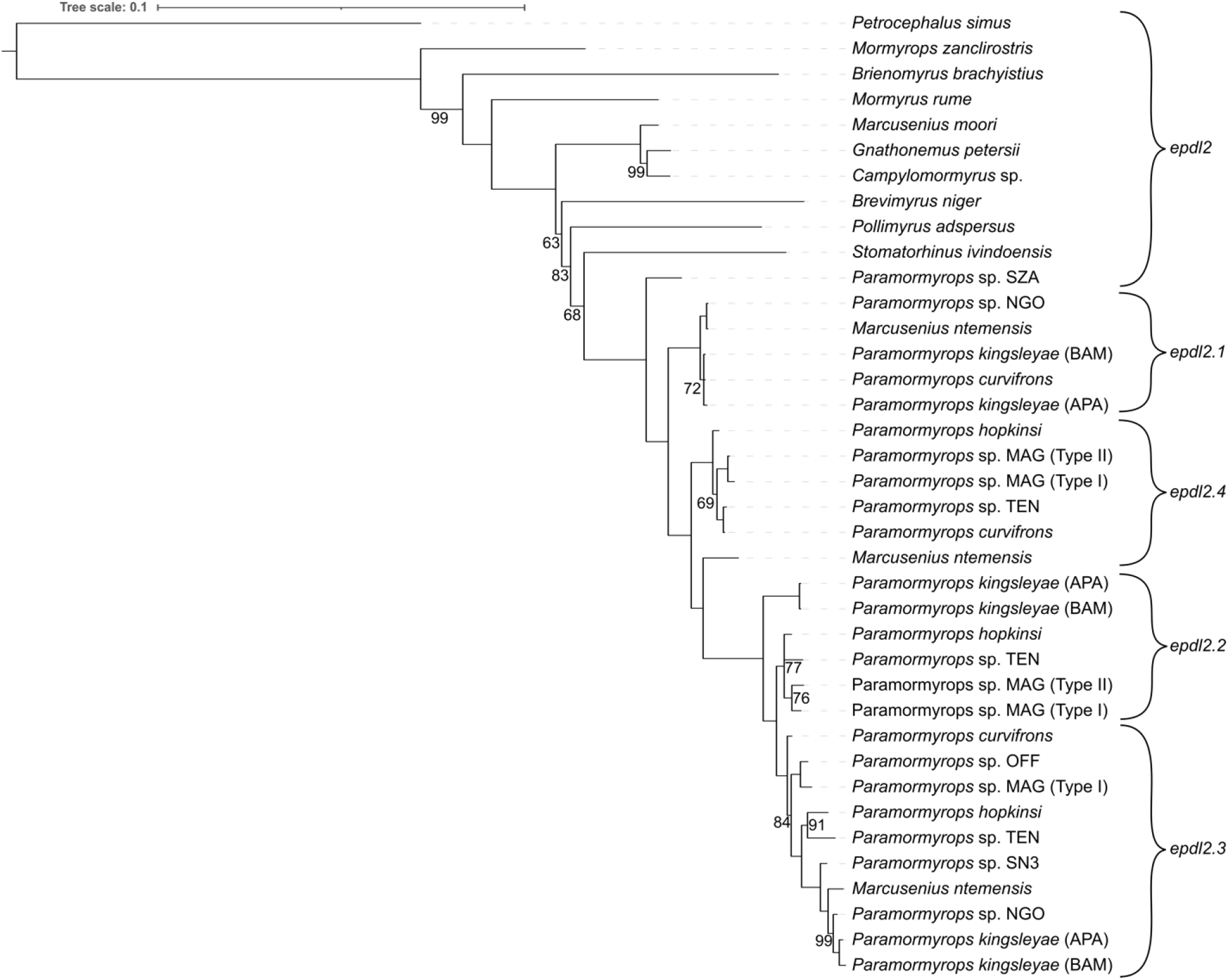
Mormyrid *epdl2* gene tree based on Bayesian inference, only posterior probabilities < 1 are shown. *epdl2* paralogs are classified based on our best hypothesis of *epdl2* duplications

We leveraged this gene tree to infer the *epdl2* duplication history. In the Mn+P clade, the earliest branch is of the sole *epdl2* gene detected in *Paramormyrops* sp. SZA, in contrast to multiple paralogs found in the rest of the clade. Therefore, we interpret that *epdl2* is not duplicated in *P*. sp. SZA. Genes orthologous to *P. kingsleyae epdl2.1* branch earliest and share the unique feature of a 130-140 bp deletion in intron 2. A second bifurcation reveals a potentially novel paralog, *epdl2.4*; whereas the remaining genes broadly sort into two clusters, orthologous to *P. kingsleyae epdl2.2* and *epdl2.3*. We note that each of the four resulting *epdl2* paralog clades contains no more than one gene from any given sample. There are two branches in this gene tree that fall outside of these paralog clusters: *M. ntemensis epdl2.4* and *P. kingsleyae epdl2.2*. We assigned these putative identities based on inspection of their CDSs compared to unambiguously assigned paralogs, and on the shortest branch distances between these sequences and the paralog clades (Fig. 4). In the case of *P. kingsleyae epdl2.2*, the existence of *P. kingsleyae epdl2.3* indicates that the sequence in question does not belong to paralog *epdl2.3*.

We summarized the distribution of *epdl2* genes across the most recent phylogenetic topology of the Mn+P clade (Peterson et al. 2022) in Fig. 5. *Epdl2.3* was detected in most species, and *epdl2.2* was consistently recorded in the clade under node A and in *P. kingsleyae*, suggesting paralog loss or critical mutations in primer-binding regions at node C and in *M. ntemensis*. The evolutionary history of paralogs *epdl2.1* and *epdl2.4* is less clear. Sequence divergence within duplicate *epdl2* genes is low, as reflected in their short branch lengths (Fig. 4). This divergence is lowest for *epdl2.1* and *epdl2.4*, and we detected both genes present in the same species in only two instances, *P. curvifrons* and *M. ntemensis* (Fig. 5). It is unlikely that these are allelic variants of the same paralog, because i) all *epdl2.1* sequences unambiguously and uniquely lack ~130 bp from intron 2, and ii) the predicted protein sequences of the two paralogs from *P. curvifrons*, and also those from *M. ntemensis*, each share a pairwise sequence identity < 94%, likely too low to be alleles from the same gene. The distribution of these two paralogs in Fig. 5 presents a scenario of multiple gene losses or failed amplifications: *epdl2.1* is missing in the clade under node A and in some species from the clade under node C; whereas *epdl2.4* is missing in *P*. kingsleyae and in some species from the clade under node C. Although our results suggest that there are four paralogs present in our data, we discuss a three-paralog scenario (where *epdl2.1* and *epdl2.4* would be the same gene) in the Discussion section.

**Fig. 5.**
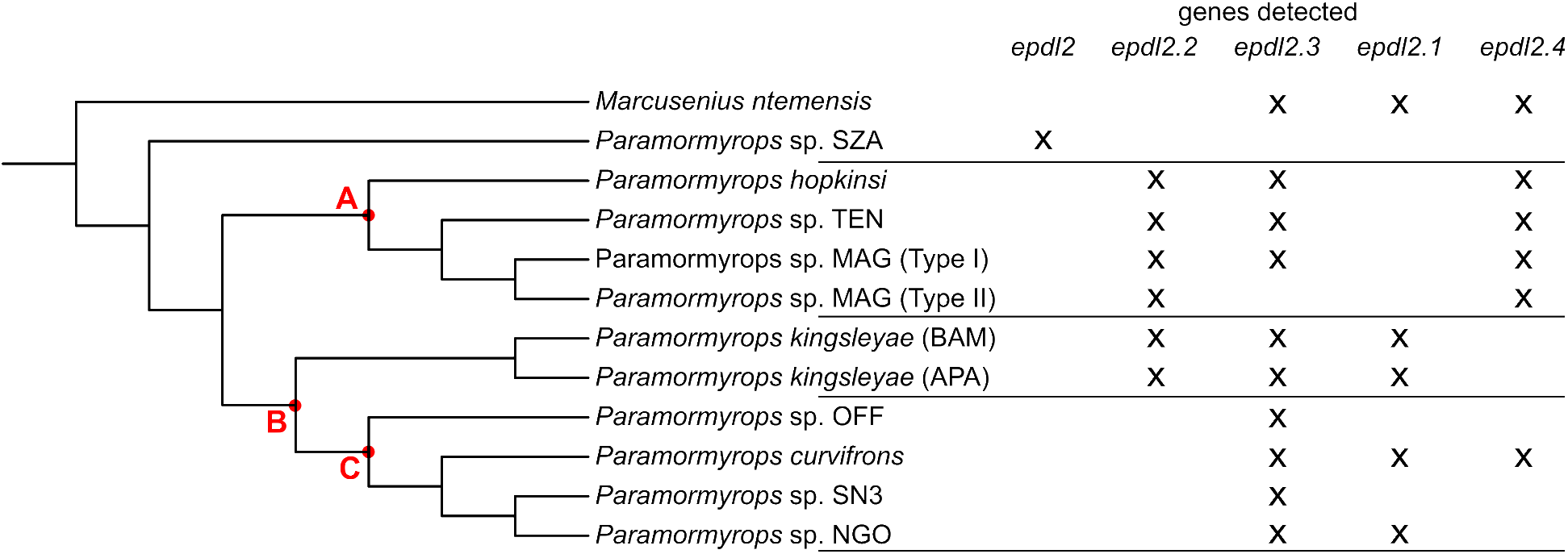
Distribution of detected *epdl2* genes across the phylogenetic topology of the *Marcusenius ntemensis* + *Paramormyrops* clade. Potential paralogs losses could have occurred at nodes A, B, or C; see main text for details.

Together, these results indicate that three rounds of *epdl2* duplication have occurred within the Mn+P clade. Since all paralogs are broadly distributed across this clade (Fig. 5), we propose that all *epdl2* duplications occurred relatively quickly and early in *Paramormyrops* evolution. If the duplications took place in an ancestor of the Mn+P clade, the non-duplicated gene may remain in *P*. sp. SZA through incomplete lineage sorting (ILS), or alternatively, the *epdl2* gene in *P*. sp. SZA may be a misclassified paralog. On the other hand, if the duplications happened in an ancestor of all the *Paramormyrops* species sequenced except *P*. sp. *SZA*, the duplicated paralogs may exist in *M. ntemensis by* way of introgression.

### Selection Tests

EOD signal evolution within the more than 20 species of *Paramormyrops* is profound (Sullivan et al. 2002; Arnegard, McIntyre, et al. 2010; Picq et al. 2020), and key features of the electric signal correlate with *epdl2* expression in mormyrid electric organs (Losilla et al. 2020, Losilla M, Gallant JR, unpublished data). These results, together with the detection of multiple *epdl2* duplications in *Paramormyrops*, motivated us to search for evidence of selection on the *epdl2* duplicates.

We did not find evidence of gene conversion in the alignment of osteoglossiform *epdl2* CDSs after running three variants of the GARD method: in all cases the models with recombination showed no improvement over the no recombination null model, Δc-AIC = 0. Thus, we deployed branch- and site-models on this alignment and the osteoglossiform *epdl2* gene tree (Fig. 6) to parse patterns of selection in our data.

**Fig. 6.**
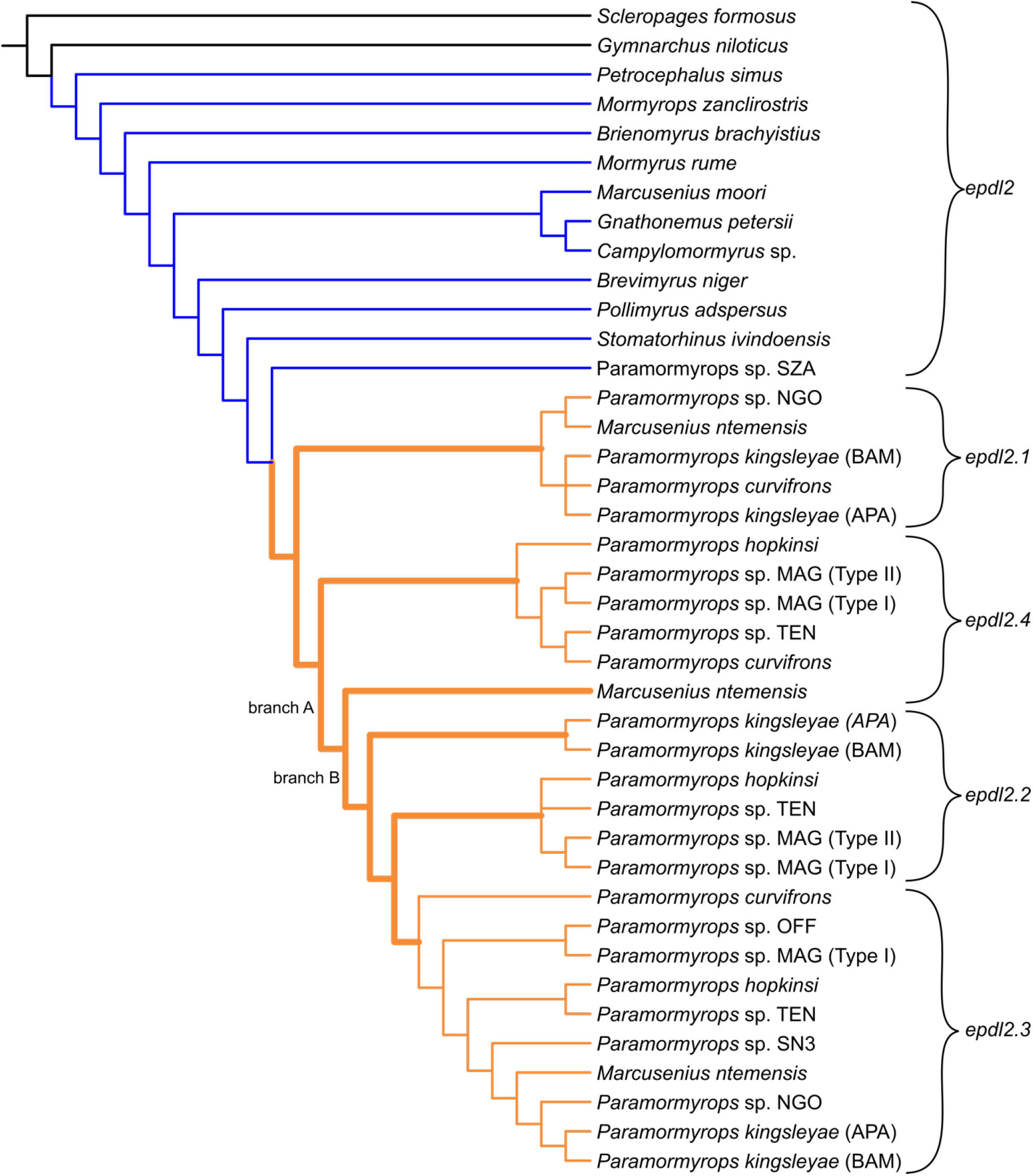
Topology of the osteoglossiform *epdl2* gene tree used in the selection tests. Mormyrid lineages with (orange) and without (blue) *epdl2* duplications are indicated. This branch partition was used in the RELAX and Contrast-FEL tests. Thick orange branches were tested for positive selection with aBSREL, and significant branches from this test are labeled A and B.

First, we compared ω values between mormyrid lineages with vs without *epdl2* duplications (Fig. 6) using the RELAX method, and found strong indications that selection intensified in the lineages with duplications (K = 6.7, LRT = 36.9, p = 1.3×10^−9^). Because this test compares ω branch values between user-defined branches and not against a value of one, it makes no claims about modes of selection. We then searched for signals of positive selection in the branches between the duplication events and the early evolution of the extant paralog genes using the aBSREL test. Conservatively, we treated the difficult-to-group branches *M. ntemensis epdl2.4* and *P. kingsleyae epdl2.2* as additional paralogs for the purpose of this test. Thus, we selected 11 branches for testing (ten were tested, aBSREL deleted one branch because it estimated it had zero length) and we found statistical support for positive selection on two of them (Fig. 6, branch A: LRT = 8.7, corrected p = 0.04; branch B: LRT = 8.8, corrected p = 0.04). These branches broadly coincide with the time of the *epdl2* duplication events.

Finally, we detected sites of interest in *epdl2*. We identified 21 *epdl2* codons that have experienced positive selection in one or more branches of the *epdl2* tree (MEME test, p < 0.1, Fig. S1). In addition, Contrast-FEL detected 36 sites evolving under different selective pressures between the mormyrid lineages with vs without *epdl2* duplications (Fig. S2, q < 0.2), with all 36 sites presenting a higher ω value in the lineages with duplications. This method does not interrogate about modes of selection, instead it contrasts site-specific ω values between user-defined branches. From these results, we obtained a list of ten sites of interest (Table S10) that i) have experienced positive selection in the *epdl2* gene tree (Fig. S1) and ii) have evolved at increased ω rates in mormyrid lineages with vs without *epdl2* duplications (Fig. S2). For each positively selected site, MEME performs an exploratory attempt to identify the branches in which it has been under positive selection. It evaluates the Empirical Bayes Factor (EBF) for observing positive selection at each branch and flags a branch as potentially under positive selection if its EBF > 100 (Murrell et al. 2012) (Table S10). Given the experimental nature of this classification, we only consider it in relation to the results from the other tests we performed (Fig. S3). Finally, we summarize the amino acid substitutions observed at these ten sites in the Epdl2 paralogs (Table S11).

### Structural predictions of Epdl2

Because we were able to identify signatures of selection at particular Epdl2 sites, we were motivated to link these sites to putative functional differences. To do so, we explored structural features using the Epdl2 amino acid sequence of *Paramormyrops* sp. SZA as a representative mormyrid Epdl2 protein. We identified a signal peptide (the first 17 amino acids) and two potential N-glycosylation sites. Also present are four conserved cysteines expected to form disulfide bonds and a highly conserved proline residue that may influence ligand binding (McDougall et al. 2018). We mapped these features and our ten sites of interest (Table S10) onto the primary structure of this reference protein (Fig. 7A) and modeled its 3D structure with I-TASSER.

**Fig. 7.**
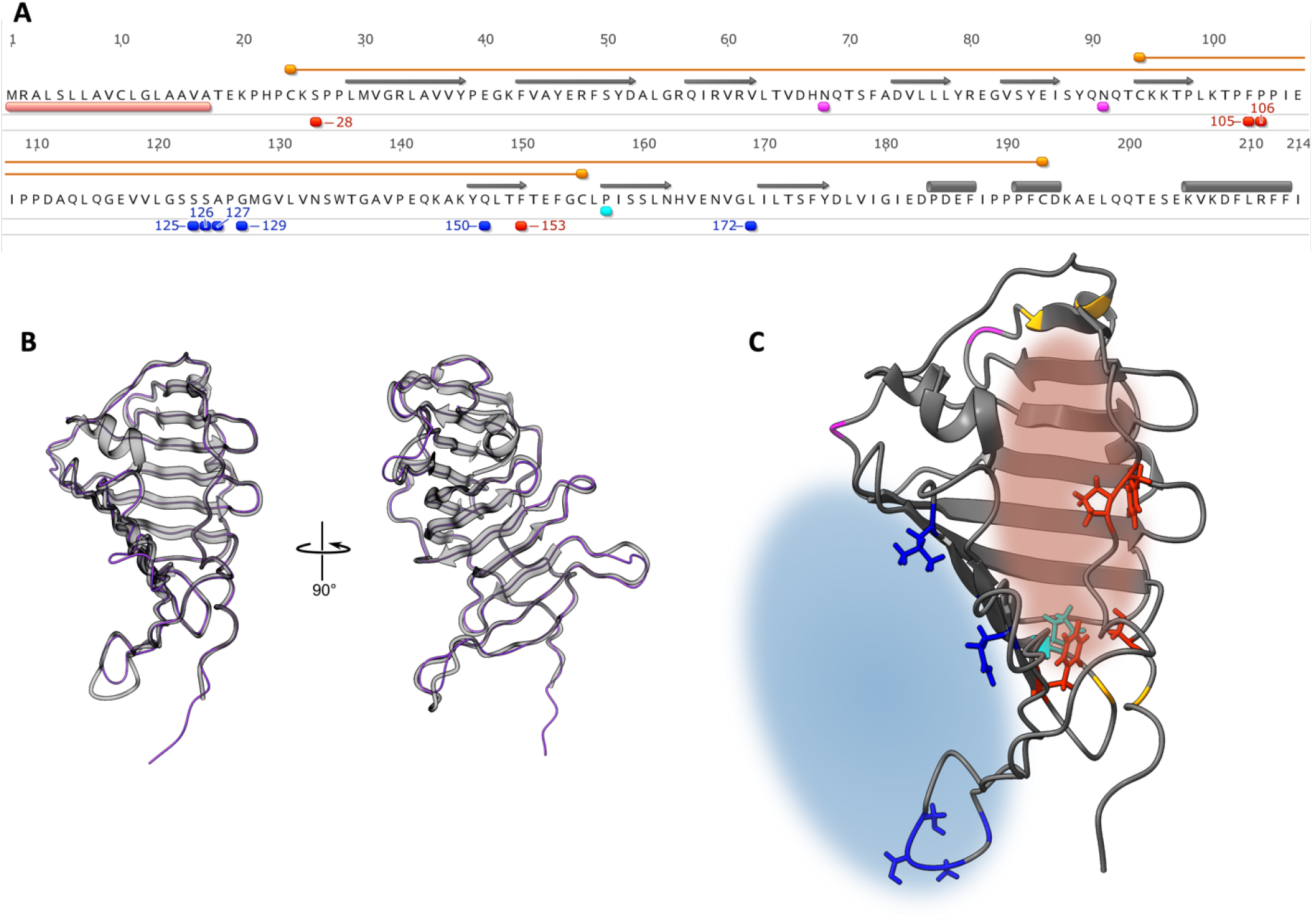
*Paramormyrops* sp. SZA Epdl2 as a representative Epdl2 protein. A) Predicted amino acid sequence, annotated with i) salient structural properties: signal peptide (pink), conserved cysteine residues (orange) forming disulfide bonds (orange lines), N-glycosylation sites (magenta), a proline residue highly conserved across EPDRs (cyan); ii) secondary structure adopted in the 3D model: β strands (gray arrows) and α helixes (gray cylinders); iii) the ten positively selected sites with increased ω rates in *epdl2* paralogs (red and blue, numbers indicate sites’ positions derived from the homologous codon alignment). B) Two views of our Epdl2 (minus the signal peptide) 3D model (gray) superimposed on its best structural analog, *Xenopus tropicalis* Epdr1 (PDB entry 6JL9, purple backbone trace). C) Our 3D model from B), showcasing structural properties and select residues as color coded in A), and predicted functional regions: ligand-binding pocket (red cloud) and dimerization surface (blue cloud). The side chains (colored sticks) of the residues depicted in red and cyan point inwards the pocket, and side chains of blue residues are oriented towards the dimerization surface.

I-TASSER produces a full-length atomic model of the user-supplied sequence, based on the topology of the best-matching known structural template, which was the crystal structure of *Xenopus tropicalis* Epdr1 (Protein Data Bank (PDB) entry 6JL9). The resulting 3D model is supported with high confidence (C-score 0.59, TM-score 0.92 to PDB hit 6JL9, Fig. 7B).

EPDRs contain a hydrophobic ligand-binding pocket, critical to its biological function (McDougall et al. 2018; Park et al. 2019; Wei et al. 2019). As expected based on these prior findings, our model predicts such a pocket (Fig. 7C, red cloud). Our 3D model prediction identified several potential ligand-binding residues and a few proposed active site residues that overlap with some of our sites of interest located in the pocket, although these predictions have low confidence.

We mapped the ten sites of interest (Table S10) on this 3D model (Fig. 7C, red and blue residues). The orientation of their side chains may be indicative to their function: Four residues (Fig. 7C, red residues) are oriented inwards the pocket and thus may participate in ligandbinding (McDougall et al. 2018; Park et al. 2019; Wei et al. 2019) and/or calcium binding (Park et al. 2019), whereas six residues (Fig. 7C, blue residues) may contribute to homodimer formation (Shashoua 1985; Hoffmann 1994), since they point outwards and belong to regions critical to Epdr1 dimerization in this protein’s solved 3D structures (Park et al. 2019; Wei et al. 2019) (Fig. 7C, blue cloud). Therefore, all our ten sites of interest likely participate in critical protein functions.

## Discussion

Weakly electric fishes have provided neat examples of duplication and subsequent neofunctionalization of ion channel genes, with key phenotypic consequences to the electric signal (Zakon et al. 2006; Arnegard, Zwickl, et al. 2010; Paul et al. 2016; Swapna et al. 2018). Here, we focus on a gene that compared to ion channels, is poorly understood. *Epdl2 is* strongly differentially expressed between mormyrids with divergent EOD signals (Losilla et al. 2020, Losilla M, Gallant JR, unpublished data) and exhibits multiple paralogous copies in one mormyrid genome. In this study, we first performed an exploratory analysis of vertebrate EPDR evolution to better understand the evolutionary context of *epdl2*. Next, we confirmed three extant, seemingly functional *epdl2* genes in *P. kingsleyae*. Then, we sequenced this gene across Mormyridae, identified three or four paralogs, and found evidence for multiple tandem duplications in an early *Paramormyrops* ancestor. We detected signs of positive selection and of increased selection rates in branches that led to *epdl2* paralogs and in sites along the gene. Finally, we generated an Epdl2 3D model and leveraged it to infer functional implications of amino acid substitutions at ten sites of interest.

### *Epdl2* and the broad evolutionary history of vertebrate EPDRs

The Genomicus EPDR tree (Fig. 2A) offers a clear hypothesis of vertebrate EPDR gene evolution: all vertebrate EPDR genes descend from an ancestral pre-vertebrate ortholog that duplicated before vertebrate diversification −potentially in one of the two vertebrate whole genome duplications (Dehal and Boore 2005; Holland and Ocampo Daza 2018). The two resulting paralogs remain widely distributed across the clade: *epdr1* is present in most if not all vertebrate species, and duplications and losses are rare, in contrast with most EPDR genes (McDougall et al. 2018). To contrast, *epdl* was lost in the water-to-land transition but has experienced duplications in bowfin and early in teleost evolution. Based on our phylogenetic analysis (Fig. 2B), we conclude that all teleost-specific EPDR genes evolved from *epdl*, and we support four teleost-specific early *epdl* paralogs: *epdl1, epdl2, epdl3*, and *epd*, plus an additional duplication in the last, resulting in *epd1* and *epd2*. Our synteny analysis showed that only *epdl1* and *epdl2* share neighboring genes with bowfin *epdl* (Fig. 2C), thus suggesting that these two teleost-specific *epdl* homologs are paralogs from a duplication event that involved multiple genes. Together, these findings suggest that *epdl2* is likely a TGD ohnolog (sensu Wolfe 2000). Its taxonomic scope is Osteoglossiformes (with further duplications in *Paramormyrops*), Otomorpha (with an additional duplication in Cyprininae), and Euteleosteomorpha, where it is present in Protacanthopterygii and in Osmeriformes, but has been lost in Neoteleostei (Fig. 2A).

While this analysis gives a broad evolutionary history of vertebrate EPDR genes, some intriguing questions remain. First, the currently known distribution of *epdl3* and *epd* throughout the main, early-branching teleost taxa is mutually exclusive, which raises the question of if these could be sequences from the same *epdl* paralog. However, we found no shared synteny between *P. kingsleyae epdl3, Danio rerio epd1*, and *Cyprinus carpio epd2*—thus making this scenario less likely. Second, our *epdl* tree rejects that a gene in *Clupea harengus* represents a different teleost *epdl* paralog, and instead clustered it with the *epdl1* sequences, although this node has the smallest posterior probability (Fig. 2B). This gene shares some neighboring genes with *epdl1* paralogs in Otocephala, thus supporting the Bayesian inference *epdl* tree topology. Third, this tree grouped *Denticeps clupeoides epdl1* with the *epd* genes. We note that there are disagreements between the Ensembl and NCBI annotations of this gene (Table S1), hence this gene may warrant closer inspection. Finally, our gene tree clusters the two osteoglossiform *epdl3* sequences with the three bowfin *epdl* sequences. The precise phylogenetic position of *epdl3* needs to be addressed in a detailed study of fish EPDR evolution.

### *Epdl2* amplicon sequencing

Our Sanger sequencing efforts confirm that there are three distinct, complete *epdl2* paralogs in tandem in *P. kingsleyae* (Fig. 3). We provide high-quality reference sequences for each gene, including complete intronic sequences and ample portions of the up- and downstream untranslated regions. Since we found no indication of additional paralogs in this region or elsewhere in the genome, we are confident that there are three *epdl2* paralogs in *P. kingsleyae*.

We were able to survey *epdl2* broadly across Mormyridae by developing a bespoke Nanopore-based amplicon sequencing approach and validated its sensitivity and accuracy with our Sanger sequenced *P. kingsleyae* genes. The combination of high-fidelity PCR, multiplexed, third-generation sequencing, and our devised bioinformatic pipeline (Fig. 1) allowed for an accurate, comprehensive sampling of mormyrid *epdl2* sequences, and would be easily extensible to other gene targets. Multiplexed amplicon Nanopore sequencing is a viable and cost-effective option to study duplicated regions across divergent taxa, and we envision our approach as a valuable tool for researchers across many study systems.

While this Nanopore-based amplicon sequencing scheme was successful, we note that this approach was limited by how conserved sequences were outside both the start and stop codons on every paralog. Importantly, the number of *epdl2* paralogs we detected for each sample (Table S2) should not be considered definitive, particularly for difficult to amplify samples from post-duplication taxa (footnotes of Tables S4 and S9). Definitive conclusions about copy number variation in *epdl2* will be greatly facilitated by future genome sequencing efforts in *M. ntemensis* and *Paramormyrops*, particularly the ~40 Kbp genomic region where these duplicates lie in tandem.

### *Epdl2* duplications occurred early in *Paramormyrops* evolution

We obtained a phylogenetically comprehensive *epdl2* gene tree that unambiguously indicates that *epdl2* duplications are not widely spread in Mormyridae, but rather confined to a recent but speciose branch centered around *Paramormyrops* and likely restricted to a large subset of these (Fig. 4). From this gene tree, we infer the existence of four *epdl2* gene copies in the Mn+P clade. However, the distribution of *epdl2.1* and *epdl2.4* across the clade (Fig. 5) compels us to consider the possibility that they are the same paralog, *epdl2.1*. Under this scenario, i) the amplicons we identified as *M. ntemensis epdl2.1* and *P. curvifrons epdl2.4* are potentially cross contamination artifacts, ii) *epdl2.1* lost ~130 bp from intron 2 at node B (Fig. 5), and iii) no *epdl2* paralog losses are predicted in *P. kingsleyae*. We did not find evidence for an *epdl2* pseudogene in this species’ genome. Regardless of which scenario is ultimately correct, the paralogs we describe likely cover the standing paralog diversity within the Mn+P clade. We sampled widely within this clade, we successfully amplified and sequenced most attempted samples, and most of the generated sequences are clearly assigned to one paralog (Fig. 4).

A more precise estimation of when the *epdl2* duplications occurred is impeded by the absence of *epdl2* sequences from taxa sister to the Mn+P clade (i.e. *Ivindomyrus, Boulengeromyrus, Cryptomyrus*; Peterson et al. 2022), and by the taxonomically incongruent placement of *M. ntemensis* inside *Paramormyrops* in the *epdl2* gene tree. The latter could be reconciled with the species tree by ILS or introgression at the *epdl2* locus after its duplications. There is strong evidence of introgression within *Paramormyrops* evolution (Sullivan et al. 2004). Alternatively, the gene we identified as a non-duplicated *epdl2* in *P*. sp. SZA could belong to one of the paralog groups. However, we note that this gene was consistently placed as the sister sequence to all *epdl2* paralogs (Fig. 4), as opposed to clustered into a paralog group, as we observe in both *P*. sp. OFF and *P*. sp SN3. In these species we detected only one *epdl2* gene (Fig. 5), yet they clearly belong to *epdl2.3* (Fig. 4).

Additional sequencing of *epdl2* paralogs in more *Paramormyrops* species and sequencing the entire *epdl2* region in taxonomically strategic species, should greatly resolve the withstanding details of the *epdl2* duplication history in this clade.

### Selection Tests

We investigated selection patterns at branch and site levels in the osteoglossiform *epdl2* gene tree (Fig. 6). The strength of selection, measured as higher ω values, unambiguously increased in the lineages with *epdl2* duplications compared to mormyrid lineages without *epdl2* duplicates (Fig. 6). Furthermore, in two lineages, presumably coinciding with the duplication events, this increase has met the detection threshold for positive selection (Fig. 6, branches A and B). This, however, is likely a conservative estimate; aBSREL’s statistical power in our dataset is limited by two technical reasons (Smith et al. 2015): i) this test requires multiple testing correction, hence we only tested a few branches; and ii) branch length directly affects statistical power, and some of the interrogated branches where we did not detect positive selection exhibit the lowest levels of divergence (paralog clusters *epdl2.1* and *epdl2.4*, Fig. 4).

Our site tests uncovered evidence of multiple sites in *epdl2* under positive selection across Mormyridae (Fig. S1), and independently, also of sites where selection has intensified in the lineages with *epdl2* duplications (Fig. S2). We focused on the ten sites common to the two sets (Table S10), because we reason that these are the sites most likely targeted by selection after the *epdl2* duplications. While we observe that some sites are supported by a substitution at a single sample (e.g. sites 105, 153) others experienced widespread, often paralog-specific substitutions (e.g. sites 106, 150) (Table S11).

Although the methodological nature of the MEME test (and similar site tests) hinders its ability to identify branch-site combinations subject to positive selection, it employs an exploratory procedure (the EBF) to suggest branches where a positively selected site may have experienced this selection (Murrell et al. 2012). These specific branch-site combinations should not be heavily relied on due to the exploratory nature of their statistical support. That said, because we screened our ten sites of interest for augmented selection in lineages with *epdl2* duplications, it is to be expected that the branches where these ten sites have experienced positive selection are post-duplication branches. Reassuringly, every branch-site pair with EBF support (Table S10) is located in lineages with duplications (Fig. S3, purple bars). Taken together, our branch and site tests converge on the conclusion that the *epdl2* paralogs have experienced significant positive selection.

### Functional Consequences of Molecular Evolution in *epdl2* paralogs

EPDR proteins are widespread across eukaryotic groups, where they participate in a diverse gamut of often lineage specific functions (Shashoua 1985; Schmidt et al. 1995; Pradel et al. 1999; Jackson et al. 2006; Zheng et al. 2006; Staats et al. 2016; Hall et al. 2017). Given their low amino acid conservation, these functions are likely accomplished through shared structural and biochemical properties: a signal peptide, N-glycosylation sites, disulfide bonds, dimerization, and a hydrophobic pocket. This suggests that comparisons between the 3D structures of Epdl2 and Epdr1 facilitate inferences about expected broad functional features of Epdl2, despite their numerous differences.

The structural annotation of *Paramormyrops* sp. SZA Epdl2 and its 3D model prediction (Fig. 7) confirm that it contains the two expected disulfide bond-forming cysteine pairs, a signal peptide on its N terminus, β strands that form two antiparallel β sheets connected by a linker region, α helixes near the C terminus (McDougall et al. 2018; Park et al. 2019; Wei et al. 2019), and two N-glycosylation sites whose position is conserved in fish EPDRs (Müller-Schmid et al. 1992; Ortí and Meyer 1996; Suárez-Castillo and García-Arrarás 2007). Although these positions are not conserved in other EPDRs including Epdr1, glycosylation in the latter may be necessary for its function (Wei et al. 2019). Our 3D model suggests that these sites are exposed in Epdl2, and therefore likely accessible to the glycosylation machinery.

The β sheets have been predicted to form a deep hydrophobic pocket in all EPDRs with potential roles in ligand-binding (McDougall et al. 2018), and its existence and functionality have been demonstrated in Epdr1 (Park et al. 2019; Wei et al. 2019). Our model showcases a striking spatial similarity with Epdr1 (Fig. 7B), in addition to a highly conserved proline residue located inside this pocket, as predicted by McDougall et al. (2018) (Fig. 7C). Every structural homology unambiguously identifies the location of this hydrophobic pocket in Epdl2 (Fig. 7C, red cloud). Epdr1’s hydrophobic pocket is similar to bacterial proteins of the LolA superfamily (Park et al. 2019; Wei et al. 2019), which participate in widespread functions through binding diverse hydrophobic ligands in their pocket (Wei et al. 2019). Sequence conservation in this region is low among EPDRs, and between Epdr1 and LolA (Wei et al. 2019). The specificity of EPDR ligands likely depends on the amino acid residues in these proteins’ hydrophobic pocket, and this apparent versatility in their binding affinities could relate to EPDRs’ involvement in a remarkable wide range of functions. It is straightforward to conceive that duplicated EPDR copies could be tailored to slightly different ligands through changes in their amino acid sequences at key positions in this pocket. Additionally, the crystal structure of Epdr1 demonstrates that two Epdr1 chains associate into a homodimer through contacts between one of the β sheets and a linker region (Park et al. 2019; Wei et al. 2019). Given the predicted structural similarity between Epdl2 and Epdr1, we expect that Epdl2 also forms homodimers mediated by its structural homologs (Fig. 7C, blue cloud).

Based on this framework, we propose putative functional consequences of the ten sites of highest interest identified in our site tests (Table S10). Four of these sites may participate in ligand binding: sites 105 and 106 are located in the pocket’s lid, and sites 28 and 153 map to the pocket’s lining, spatially close to the conserved proline site. All four residues have their side chains oriented towards the inside of the pocket (Fig. 7C, red residues). We presume that the other six sites are consequential for dimer formation: sites 150 and 172 belong in the β sheet that participates in this process, and sites 125, 126, 127, and 129 are located within the linker region involved in contacts between the monomeric subunits. The side chains of all six residues point towards the dimerization surface (Fig. 7C, blue residues). We note that a necessary step towards efficient subfunctionalization in duplicated Epdl2 proteins is the correct pairing of the Epdl2 paralog chains into homodimers, which is likely to be achieved by paralog-specific amino acid changes in the dimerization surface.

Collectively, our results suggest selection-driven diversification in amino acid residues that coherently affects protein function. We emphasize that the procedure we used to identify signatures of selection relies on coding sequences and is therefore uninformed by protein structures, and the 3D model prediction is not aware of our sites of interest.

## Conclusions

In this analysis, we sequenced and annotated the three *epdl2* paralogs present in *P. kingsleyae* and developed a novel, Nanopore-based, sequencing and bioinformatics approach to analyze amplicons from duplicated targets, which yielded *epdl2* sequences from 20 mormyrid taxa. We propose that as many as four *epdl2* paralogs resulted from tandem duplication events in an early *Paramormyrops* ancestor, and we demonstrate that these *epdl2* paralogs, and specific sites in them, have experienced increased selection rates and have been targets of positive selection. Finally, we identify that these specific sites are located in protein regions relevant to ligand binding and homodimer formation. Presently, the specific functional role of Epdl2 in mormyrid electrocytes is unknown. The homodimeric configuration of Epdr1, and hence the expected arrangement in Epdl2, contains both hydrophobic pockets next to each other, open towards a relatively flat surface on the dimer. It has been proposed that this allows membrane binding “with an extensive contact surface for the binding and possible extraction and solubilization of target lipids” (Wei et al. 2019, p. 8). We hypothesize based on these proposed functions that Epdl2 may be related to the shaping/maintenance of the electrocyte’s plasma membrane via selectively altering its lipid components. Selective changes to membrane lipids could change their composition and distribution, and therefore influence the membrane’s biochemical properties like shape, curvature, electrical charges, fluidity, and local protein composition. Given the well-established connection between the electrocytes’ membrane traits and the resulting EOD (Bennett and Grundfest 1961; Szabo 1961; Bennett 1971; Hopkins 1999; Gallant et al. 2011; Carlson and Gallant 2013), increased plasticity and control of this membrane’s biophysical properties could facilitate EOD divergence, which may contribute to species diversification.

## Acknowledgements

Several of the samples utilized in this paper were captured by ‘Equipe Bafoule’ consisting of Dr. Sophie Picq, Dr. Lauren Koenig, Hans Mipounga, Nestor Ngoya, Franck Nzigou, and staff at Centre National de la Recherche Scientifique et Technologique (CENAREST) in Libreville, Gabon. Dr. John Sullivan provided key assistance in identifying species from the Gabon 2019 expedition. Additional samples were provided by Dr. Kerri Ackerley, Dr. Bruce Carlson, and Erika Schumacher. We would also like to acknowledge Drs. Stacy Pirro and Rose Peterson for providing genomic scaffolds from multiple mormyrid species to aid in primer design in the early stages of this project, and Dr. Ingo Braasch for helpful advice and feedback on earlier versions of this manuscript. This work was supported through computational resources and services provided by the Institute for Cyber-Enabled Research at Michigan State University, the National Science Foundation (grant numbers 1455405, 1856243), and a Michigan State University College of Natural Science Barnett Rosenberg Endowed Research Assistantship to ML.

## Notes

### Competing Interest Statement

The authors have declared no competing interest.

### Summary of Updates

The remaining GenBank/NCBI accession numbers have been added. This action produced a few, minor changes to the main text and the supplemental files

## References

Abascal F, Zardoya R, Telford MJ. 2010. TranslatorX: Multiple alignment of nucleotide sequences guided by amino acid translations. Nucleic Acids Research 38:7–13.

Almagro Armenteros JJ, Tsirigos KD, Sønderby CK, Petersen TN, Winther O, Brunak S, von Heijne G, Nielsen H. 2019. SignalP 5.0 improves signal peptide predictions using deep neural networks. Nature Biotechnology 37:420–423.

Alves-Gomes J, Hopkins CD. 1997. Molecular insights into the phylogeny of mormyriform fishes and the evolution of their electric organs. Brain Behav Evol 49:324–350.

Anderson RP, Roth JR. 1977. Tandem genetic duplications in phage and bacteria. Annual Review of Microbiology [Internet] 31:473–505. Available from: https://doi.org/10.1146/annurev.mi.31.100177.002353

Arnegard ME, Bogdanowicz SM, Hopkins CD. 2005. Multiple cases of striking genetic similarity between alternate electric fish signal morphs in sympatry. Evolution 59:324–343.

Arnegard ME, Hopkins CD. 2003. Electric signal variation among seven blunt-snouted Brienomyrus species (Teleostei: Mormyridae) from a riverine species flock in Gabon, Central Africa. Environmental Biology of Fishes 67:321–339.

Arnegard ME, McIntyre PB, Harmon LJ, Zelditch ML, Crampton WGR, Davis JK, Sullivan JP, Lavoué S, Hopkins CD. 2010. Sexual signal evolution outpaces ecological divergence during electric fish species radiation. Am Nat 176:335–356.

Arnegard ME, Zwickl DJ, Lu Y, Zakon HH. 2010. Old gene duplication facilitates origin and diversification of an innovative communication system--twice. Proc Natl Acad Sci U S A 107:22172–22177.

Bailey JA, Gu Z, Clark RA, Reinert K, Samonte R v., Schwartz S, Adams MD, Myers EW, Li PW, Eichler EE. 2002. Recent Segmental Duplications in the Human Genome. Science (1979) 297:1003–1007.

Bass AH. 1986. Species differences in electric organs of mormyrids: Substrates for species□ typical electric organ discharge waveforms. Journal of Comparative Neurology 244:313–330.

Bennett MVL. 1971. Electric Organs. In: Hoar WS, Randall DJ, editors. Fish Physiology. Vol. 5. Fish Physiology. London: Academic Press. p. 347–491.

Bennett MVL, Grundfest H. 1961. Studies on the morphology and electrophysiology of electric organs. III. Electrophysiology of electric organs in mormyrids. In: Chagas C, de Carvalho A, editors. Bioelectrogenesis. Amsterdam: Elsevier. p. 113–135.

Carlson BA, Gallant JR. 2013. From sequence to spike to spark: evo-devo-neuroethology of electric communication in mormyrid fishes. J Neurogenet 27:106–129.

Carlson BA, Hasan SM, Hollmann M, Miller DB, Harmon LJ, Arnegard ME. 2011. Brain evolution triggers increased diversification of electric fishes. Science (1979) 332:583–586.

Chapal M, Mintzer S, Brodsky S, Carmi M, Barkai N. 2019. Resolving noise-control conflict by gene duplication. PLoS Biology 17.

Conant GC, Wolfe KH. 2008. Turning a hobby into a job: How duplicated genes find new functions. Nature Reviews Genetics 9:938–950.

De Coster W, D’Hert S, Schultz DT, Cruts M, Van Broeckhoven C. 2018. NanoPack: Visualizing and processing long-read sequencing data. Bioinformatics 34:2666–2669.

Dehal P, Boore JL. 2005. Two Rounds of Whole Genome Duplication in the Ancestral Vertebrate. PLoS Biology 3:e314.

Edgar RC. 2004. MUSCLE: Multiple sequence alignment with high accuracy and high throughput. Nucleic Acids Research 32:1792–1797.

von der Emde G, Amey M, Engelmann J, Fetz S, Folde C, Hollmann M, Metzen M, Pusch R. 2008. Active electrolocation in Gnathonemus petersii: Behaviour, sensory performance, and receptor systems. Journal of Physiology - Paris 102:279–290.

Gallant JR, Arnegard ME, Sullivan JP, Carlson BA, Hopkins CD. 2011. Signal variation and its morphological correlates in Paramormyrops kingsleyae provide insight into the evolution of electrogenic signal diversity in mormyrid electric fish. Journal of Comparative Physiology A [Internet] 197:799–817. Available from: http://link.springer.com/article/10.1007/s00359-011-0643-8/fulltext.html

Gallant JR, Losilla M, Tomlinson C, Warren WC. 2017. The Genome and Adult Somatic Transcriptome of the Mormyrid Electric Fish Paramormyrops kingsleyae. Genome Biology and Evolution [Internet] 9:3525–3530. Available from: https://academic.oup.com/gbe/article/9/12/3525/4731783

Ganss B, Hoffmann W. 2009. Calcium-induced conformational transition of trout ependymins monitored by tryptophan fluorescence. The Open Biochemistry Journal 3:14–17.

Guindon S, Dufayard JF, Lefort V, Anisimova M, Hordijk W, Gascuel O. 2010. New algorithms and methods to estimate maximum-likelihood phylogenies: Assessing the performance of PhyML 3.0. Systematic Biology 59:307–321.

Gupta R, Brunak S. 2002. Prediction of glycosylation across the human proteome and the correlation to protein function. Pacific Symposium on Biocomputing 7:310–322.

Hall MR, Kocot KM, Baughman KW, Fernandez-Valverde SL, Gauthier MEA, Hatleberg WL, Krishnan A, McDougall C, Motti CA, Shoguchi E, et al. 2017. The crown-of-thorns starfish genome as a guide for biocontrol of this coral reef pest. Nature [Internet] 544:231–234. Available from: http://dx.doi.org/10.1038/nature22033

Hoffmann W. 1994. Ependymins and their potential role in neuroplasticity and regeneration: calcium-binding meningeal glycoproteins of the cerebrospinal fluid and extracellular matrix. Int J Biochem 26:607–619.

Holland LZ, Ocampo Daza D. 2018. A new look at an old question: when did the second whole genome duplication occur in vertebrate evolution? Genome Biology 19:209.

Hopkins CD. 1981. On the diversity of electric signals in a community of mormyrid electric fish in West Africa. American Zoologist 21:211–222.

Hopkins CD. 1999. Signal evolution in electric communication. In: Hauser M, Konishi M, editors. The design of animal communication. Cambridge, MA: MIT Press. p. 461–491.

Huelsenbeck JP, Ronquist F. 2001. MRBAYES: Bayesian inference of phylogenetic trees. Bioinformatics 17:754–755.

Jackson DJ, McDougall C, Green K, Simpson F, Wörheide G, Degnan BM. 2006. A rapidly evolving secretome builds and patterns a sea shell. BMC Biology 4:1–10.

Koren S, Walenz BP, Berlin K, Miller JR, Bergman NH, Phillippy AM. 2017. Canu: scalable and accurate long-read assembly via adaptive k-mer weighting and repeat separation. Genome Research 27:722–736.

Kosakovsky Pond SL, Frost SDW, Muse S V. 2005. HyPhy: hypothesis testing using phylogenies. Bioinformatics 21:676–679.

Kosakovsky Pond SL, Poon AFY, Velazquez R, Weaver S, Hepler NL, Murrell B, Shank SD, Magalis BR, Bouvier D, Nekrutenko A, et al. 2020. HyPhy 2.5 - A Customizable Platform for Evolutionary Hypothesis Testing Using Phylogenies. Molecular Biology and Evolution 37:295–299.

Kosakovsky Pond SL, Posada D, Gravenor MB, Woelk CH, Frost SDW. 2006. Automated phylogenetic detection of recombination using a genetic algorithm. Molecular Biology and Evolution 23:1891–1901.

Kosakovsky Pond SL, Wisotsky SR, Escalante A, Magalis BR, Weaver S. 2021. Contrast-FEL-A Test for Differences in Selective Pressures at Individual Sites among Clades and Sets of Branches. Mol Biol Evol 38:1184–1198.

Kramer B. 1974. Electric organ discharge interaction during interspecific agonistic behaviour in freely swimming mormyrid fish. Journal of Comparative Physiology 93:203–235.

Lavoué S, Arnegard ME, Sullivan JP, Hopkins CD. 2008. Petrocephalus of Odzala offer insights into evolutionary patterns of signal diversification in the Mormyridae, a family of weakly electrogenic fishes from Africa. Journal of Physiology Paris 102:322–339.

Lavoué S, Sullivan JP, Hopkins CD. 2003. Phylogenetic utility of the first two introns of the S7 ribosomal protein gene in African electric fishes (Mormyroidea: Teleostei) and congruence with other molecular markers. Biological Journal of the Linnean Society [Internet] 78:273–292. Available from: https://academic.oup.com/biolinnean/article-lookup/doi/10.1046/j.1095-8312.2003.00170.x

Lefort V, Longueville JE, Gascuel O. 2017. SMS: Smart Model Selection in PhyML. Mol Biol Evol 34:2422–2424.

Lemoine F, Correia D, Lefort V, Doppelt-Azeroual O, Mareuil F, Cohen-Boulakia S, Gascuel O. 2019. NGPhylogeny.fr: new generation phylogenetic services for non-specialists. Nucleic Acids Research [Internet] 47:W260–W265. Available from: https://doi.org/10.1093/nar/gkz303

Li W, Godzik A. 2006. Cd-hit: A fast program for clustering and comparing large sets of protein or nucleotide sequences. Bioinformatics 22:1658–1659.

Lipinski KJ, Farslow JC, Fitzpatrick KA, Lynch M, Katju V, Bergthorsson U. 2011. High Spontaneous Rate of Gene Duplication in Caenorhabditis elegans. Current Biology 21:306–310.

Lissmann HW, Machin KE. 1958. The mechanism of object location in Gymnarchus niloticus and similar fish. Journal of Experimental Biology 35:451–486.

Losilla M, Luecke DM, Gallant JR. 2020. The transcriptional correlates of divergent electric organ discharges in Paramormyrops electric fish. BMC Evolutionary Biology [Internet] 20:6. Available from: https://bmcevolbiol.biomedcentral.com/articles/10.1186/s12862-019-1572-3

Löytynoja A, Goldman N. 2005. An algorithm for progressive multiple alignment of sequences with insertions. Proc Natl Acad Sci U S A 102:10557–10562.

Lynch M, Conery JS. 2000. The Evolutionary Fate and Consequences of Duplicate Genes. Science (1979) 290:1151–1155.

Lynch M, Sung W, Morris K, Coffey N, Landry CR, Dopman EB, Dickinson WJ, Okamoto K, Kulkarni S, Hartl DL, et al. 2008. A genome-wide view of the spectrum of spontaneous mutations in yeast. Proceedings of the National Academy of Sciences 105:9272–9277.

Magadum S, Banerjee U, Murugan P, Gangapur D, Ravikesavan R. 2013. Gene duplication as a major force in evolution. Journal of Genetics 92:155–161.

McDougall C, Hammond MJ, Dailey SC, Somorjai IML, Cummins SF, Degnan BM. 2018. The evolution of ependymin-related proteins. BMC Evolutionary Biology [Internet] 18:182. Available from: https://bmcevolbiol.biomedcentral.com/articles/10.1186/s12862-018-1306-y

Möhres FP. 1957. Elektrische Entladungen im Dienste der Revierabgrenzung bei Fischen. Naturwissenschaften 44:431–432.

Müller-Schmid A, Rinder H, Lottspeich F, Gertzen E-M, Hoffmann W. 1992. Ependymins from the cerebrospinal fluid of salmonid fish: gene structure and molecular characterization. Gene 118:189–196.

Murrell B, Wertheim JO, Moola S, Weighill T, Scheffler K, Kosakovsky Pond SL. 2012. Detecting individual sites subject to episodic diversifying selection. PLoS Genetics [Internet] 8:e1002764. Available from: https://journals.plos.org/plosgenetics/article?id=10.1371/journal.pgen.1002764

Nguyen NTT, Vincens P, Dufayard JF, Roest Crollius H, Louis A. 2022. Genomicus in 2022: comparative tools for thousands of genomes and reconstructed ancestors. Nucleic Acids Research [Internet] 50:D1025–D1031. Available from: https://doi.org/10.1093/nar/gkab1091

Ohno S. 1970. Evolution by gene duplication. Springer-Verlag

Ortí G, Meyer A. 1996. Molecular evolution of ependymin and the phylogenetic resolution of early divergences among euteleost fishes. Mol Biol Evol [Internet] 13:556–573. Available from: http://www.ncbi.nlm.nih.gov/pubmed/8882499

Parey E, Louis A, Montfort J, Guiguen Y, Crollius HR, Berthelot C. 2022. Genome-wide mechanisms of rediploidization in paleopolyploid teleosts revealed by a high-resolution comparative atlas across 74 fishes. bioRxiv [Internet]:2022.01.13.476171. Available from: http://biorxiv.org/content/early/2022/05/04/2022.01.13.476171.abstract

Park JK, Kim KY, Sim YW, Kim YI, Kim JK, Lee C, Han J, Kim CU, Lee JE, Park SY. 2019. Structures of three ependymin-related proteins suggest their function as a hydrophobic molecule binder. IUCrJ 6:729–739.

Paul C, Kirschbaum F, Mamonekene V, Tiedemann R. 2016. Evidence for Non-neutral Evolution in a Sodium Channel Gene in African Weakly Electric Fish (Campylomormyrus, Mormyridae). Journal of Molecular Evolution [Internet] 83:61–77. Available from: http://link.springer.com/10.1007/s00239-016-9754-8

Peterson RD, Sullivan JP, Hopkins CD, Santaquiteria A, Dillman CB, Pirro S, Betancur-R R, Arcila D, Hughes LC, Ortí G. 2022. Phylogenomics of Bony-Tongue Fishes (Osteoglossomorpha) Shed Light on the Craniofacial Evolution and Biogeography of the Weakly Electric Clade (Mormyridae). Systematic Biology.

Pettersen EF, Goddard TD, Huang CC, Meng EC, Couch GS, Croll TI, Morris JH, Ferrin TE. 2021. UCSF ChimeraX: Structure visualization for researchers, educators, and developers. Protein Science 30:70–82.

Picq S, Sperling J, Cheng CJ, Carlson BA, Gallant JR. 2020. Genetic drift does not sufficiently explain patterns of electric signal variation among populations of the mormyrid electric fish Paramormyrops kingsleyae. Evolution (N Y):1–25.

Pradel G, Schachner M, Schmidt R. 1999. Inhibition of memory consolidation by antibodies against cell adhesion molecules after active avoidance conditioning in zebrafish. Journal of Neurobiology 39:197–206.

Rabosky DL, Santini F, Eastman J, Smith SA, Sidlauskas B, Chang J, Alfaro ME. 2013. Rates of speciation and morphological evolution are correlated across the largest vertebrate radiation. Nature Communications 4:1–8.

Roy A, Kucukural A, Zhang Y. 2010. I-TASSER: A unified platform for automated protein structure and function prediction. Nature Protocols 5:725–738.

Schmidt R, Brysch W, Rother S, Schlingensiepen K. 1995. Inhibition of Memory Consolidation After Active Avoidance Conditioning by Antisense Intervention with Ependymin Gene Expression. Journal of Neurochemistry 65:1465–1471.

Schwarz H, Miiller-schmid A, Hoffmann W. 1993. Ultrastructural localization of ependymins in the endomeninx of the brain of the rainbow trout: possible association with collagen fibrils of the extracellular matrix. Cell Tissue:417–425.

Shashoua VE. 1985. The role of brain extracellular proteins in neuroplasticity and learning. Cell Mol Neurobiol [Internet] 5:183–207. Available from: http://www.ncbi.nlm.nih.gov/pubmed/4028066

Shen W, Le S, Li Y, Hu F. 2016. SeqKit: A cross-platform and ultrafast toolkit for FASTA/Q file manipulation. PLoS ONE 11:1–10.

Smith MD, Wertheim JO, Weaver S, Murrell B, Scheffler K, Kosakovsky Pond SL. 2015. Less is more: An adaptive branch-site random effects model for efficient detection of episodic diversifying selection. Molecular Biology and Evolution 32:1342–1353.

Staats KA, Wu T, Gan BS, O’Gorman DB, Ophoff RA. 2016. Dupuytren’s disease susceptibility gene, EPDR1, is involved in myofibroblast contractility. Journal of Dermatological Science [Internet] 83:131–137. Available from: http://www.ncbi.nlm.nih.gov/pubmed/27245865

Suárez-Castillo EC, García-Arrarás JE. 2007. Molecular evolution of the ependymin protein family: a necessary update. BMC Evol Biol [Internet] 7:23. Available from: http://www.biomedcentral.com/1471-2148/7/23

Sullivan JP, Lavoué S, Arnegard ME, Hopkins CD. 2004. AFLPs resolve phylogeny and reveal mitochondrial introgression within a species flock of African electric fish (Mormyroidea: Teleostei). Evolution [Internet] 58:825–841. Available from: http://www.ncbi.nlm.nih.gov/pubmed/15154558

Sullivan JP, Lavoué S, Hopkins CD. 2000. Molecular systematics of the African electric fishes (Mormyroidea: teleostei) and a model for the evolution of their electric organs. J Exp Biol 203:665–683.

Sullivan JP, Lavoué S, Hopkins CD. 2002. Discovery and phylogenetic analysis of a riverine species flock of African electric fishes (Mormyridae: Teleostei). Evolution (N Y) [Internet] 56:597–616. Available from: http://www.bioone.org/doi/abs/10.1554/0014-3820(2002)056%5B0597:DAPAOA%5D2.0.CO%3B2

Swapna I, Ghezzi A, York JM, Markham MR, Halling DB, Lu Y, Gallant JR, Zakon HH. 2018. Electrostatic Tuning of a Potassium Channel in Electric Fish. Current Biology [Internet] 28:2094–2102.e5. Available from: https://www.sciencedirect.com/science/article/pii/S0960982218306146

Szabo T. 1961. Les Organes Electriques des Mormyrides. In: Chagas C, de Carvalho A, editors. Bioelectrogenesis. New York: Elsevier. p. 20–24.

Taylor JS, Raes J. 2004. Duplication and divergence: The evolution of new genes and old ideas. Annual Review of Genetics 38:615–643.

Weaver S, Shank SD, Spielman SJ, Li M, Muse S V., Kosakovsky Pond SL. 2018. Datamonkey 2.0: A modern web application for characterizing selective and other evolutionary processes. Molecular Biology and Evolution 35:773–777.

Wei Y, Xiong ZJ, Li J, Zou C, Cairo CW, Klassen JS, Privé GG. 2019. Crystal structures of human lysosomal EPDR1 reveal homology with the superfamily of bacterial lipoprotein transporters. Communications Biology [Internet] 2:1–13. Available from: http://dx.doi.org/10.1038/s42003-018-0262-9

Wertheim JO, Murrell B, Smith MD, Kosakovsky Pond SL, Scheffler K. 2015. RELAX: Detecting relaxed selection in a phylogenetic framework. Molecular Biology and Evolution 32:820–832.

Wolfe K. 2000. Robustness—it’s not where you think it is. Nature Genetics 25:3–4.

Yang J, Yan R, Roy A, Xu D, Poisson J, Zhang Y. 2015. The I-TASSER suite: Protein structure and function prediction. Nature Methods 12:7–8.

Yang J, Zhang Y. 2015. I-TASSER server: New development for protein structure and function predictions. Nucleic Acids Research 43:W174–W181.

Zakon HH, Lu Y, Zwickl DJ, Hillis DM. 2006. Sodium channel genes and the evolution of diversity in communication signals of electric fishes: Convergent molecular evolution. Proceedings of the National Academy of Sciences [Internet] 103:3675–3680. Available from: http://www.pnas.org/cgi/doi/10.1073/pnas.0600160103

Zhang J. 2003. Evolution by gene duplication: An update. Trends in Ecology and Evolution 18:292–298.

Zheng FX, Sun XQ, Fang BH, Hong XG, Zhang JX. 2006. Comparative analysis of genes expressed in regenerating intestine and non-eviscerated intestine of Apostichopus japonicus Selenka (Aspidochirotida: Stichopodidae) and cloning of ependymin gene. Hydrobiologia 571:109–122.

