## Supplemental Tables and Figures for "Molecular evolution of the ependymin-related gene *epdl2* in African weakly electric fish"

Table S1. Details of the sequences used in the EPDR phylogenetic analysis

| Species | Gene | Code | | | Source | Notes |
|---|---|---|---|---|---|---|
| | | gene | transcript | protein | | |
| *Danio rerio* | *epd1* | ENSDARG00000103498 | ENSDART00000171617.2 | ENSDARP00000134034.1 | Ensembl | |
| *Denticeps clupeoides* | *epd1* | ENSDCDG00000020214 | ENSDCDT00000036790 | ENSDCDP00000027731.1 | Ensembl | |
| *Salmo salar* | *epd1* | ENSSSAG00000073991 | ENSSSAT00000133331.1 | ENSSSAP00000098870.1 | Ensembl | |
| *Takifugu rubripes* | *epd1* | ENSTRUG00000014952 | ENSTRUT00000038357.3 | ENSTRUP00000038220.2 | Ensembl | |
| *Cyprinus carpio* | *epd2* | ENSCCRG00000012904 | ENSCCRT00000025647.1 | ENSCCRP00000023616.1 | Ensembl | |
| *Pygocentrus nattereri* | *epd2* | LOC108432735 | XM_017706796.2 | XP_017562285.1 | NCBI | Splicing disagreement with Ensembl ENSPNAG00000025130 |
| *Takifugu rubripes* | *epd2* | ENSTRUG00000008083 | ENSTRUT00000020238.3 | ENSTRUP00000020154.3 | Ensembl | Manually curated due to missing exons[b] |
| *Amia calva* | *epdl* | AMCG00015468 | AMCT00015468 | AMCP00015468 | NCBI GCA_017591485.1, https://github.com/AndrewWT/AmiaGenomics | Manually curated due to missing exons[b] |
| *Amia calva* | *epdl* | AMCG00015463 | AMCT00015463 | AMCP00015463 | | Manually curated due to missing exons[b] |
| *Amia calva* | *epdl* | AMCG00015472 | AMCT00015472 | AMCP00015472 | | |
| *Latimeria chalumnae* | *epdl* | ENSLACG00000018325.2 | ENSLACT00000025567.1 | ENSLACP00000023190.1 | Ensembl | |
| *Protopterus annectens* | *epdl* | LOC122805608 | XM_044075849.1 | XP_043931784.1 | NCBI | |
| *Rhincodon typus* | *epdl* | LOC109918485 | XM_020518419.1 | XP_020374008.1 | NCBI | |
| *Rhincodon typus* | *epdl* | LOC109936335 | XM_020534949.1 | XP_020390538.1 | NCBI | Manually curated due to missing exons[b] |
| *Clupea harengus* | *epdl1*[a] | ENSCHAG00000013904 | ENSCHAT00000031403.1 | ENSCHAP00000028894.1 | Ensembl | Annotated as 3 genes in NCBI and 1 gene with 7 transcripts in Ensembl. We suspect there |

| | | | | | | |
|---|---|---|---|---|---|---|
| | | | | | | are two genes in tandem |
| *Danio rerio* | *epdl1* | ENSDARG00000076386 | ENSDART00000111531.4 | ENSDARP00000101021.2 | Ensembl | |
| *Denticeps clupeoides* | *epdl1* | LOC114788914 | XM_028977867.1 | XP_028833700 | NCBI | Splicing disagreement with Ensembl ENSDCDG00000018927 |
| *Paramormyrops kingsleyae* | *epdl1* | ENSPKIG00000021470 | ENSPKIT00000028285.1 | ENSPKIP00000004303.1 | Ensembl | |
| *Salmo salar* | *epdl1* | ENSSSAG00000002722 | ENSSSAT00000005774.1 | ENSSSAP00000005381.1 | Ensembl | |
| *Scleropages formosus* | *epdl1* | ENSSFOG00015021928 | ENSSFOT00015034799.1 | ENSSFOP00015034421 | Ensembl | This gene was retired from Ensembl with no successors. However, blast search and synteny inspection suggest this is *epdl1* |
| *Takifugu rubripes* | *epdl1* | ENSTRUG00000012059 | ENSTRUT00000030623.3 | ENSTRUP00000030506.2 | Ensembl | |
| *Danio rerio* | *epdl2* | ENSDARG00000055539.7 | ENSDART00000077910.7 | ENSDARP00000072376.5 | Ensembl | |
| *Denticeps clupeoides* | *epdl2* | ENSDCDG00000027589 | ENSDCDT00000054119 | ENSDCDP00000043043.1 | Ensembl | |
| *Paramormyrops kingsleyae* | *epdl2* | *epdl2.1* | *epdl2.1* | *epdl2.1* | This work. GenBank accession number ON863825 | Sanger-sequenced and manually curated. There were sequencing errors in coding homopolymers |
| *Protosalanx hyalocranius* | *epdl2* | LS_GLEAN_10003579 | LS_GLEAN_10003579 | LS_GLEAN_10003579 | http://gigadb.org/dataset/100262 | Manually curated due to missing exons[b] |
| *Salmo salar* | *epdl2* | ENSSSAG00000010021 | ENSSSAT00000022032.1 | ENSSSAP00000020584.1 | Ensembl | |
| *Scleropages formosus* | *epdl2* | ENSSFOG00015021850 | ENSSFOT00015034652.2 | ENSSFOP00015034274.1 | Ensembl | |
| *Brienomyrus brachyistius* | *epdl3* | LOC125720270 | XM_048995478.1 | XP_048851435.1 | NCBI | Manually curated due to missing exons[b] |
| *Paramormyrops kingsleyae* | *epdl3* | ENSPKIG00000018439 | ENSPKIT00000023217.1 | ENSPKIP00000011278.1 | Ensembl | |

[a]Classified as a unique *epdl* paralog in the Genomicus gene tree (main text Fig. 2A), but reclassified as *epdl1* by the phylogenetic analysis (main text Fig. 2B)
[b]Sequence is supplied in additional file 1

Table S2. Details of the *epdl2* genes found in every species studied

<table>
<tr><th>species</th><th>source of epdl2 sequence(s)</th><th>epdl2 genes</th><th>gene size (bp)[a]</th><th>GenBank accession numbers</th></tr>
<tr><td>Brevimyrus niger</td><td>DNA extraction, PCR amplification & multiplexed amplicon sequencing with ONT</td><td>epdl2</td><td>1823</td><td>ON863837</td></tr>
<tr><td>Brienomyrus brachyistius</td><td>genome assembly available (NCBI Gene ID 125720044)</td><td>epdl2</td><td>2166</td><td>-</td></tr>
<tr><td>Campylomormyrus sp.</td><td>DNA extraction, PCR amplification & multiplexed amplicon sequencing with ONT</td><td>epdl2</td><td>1801</td><td>ON863835</td></tr>
<tr><td>Gnathonemus petersii</td><td>DNA extraction, PCR amplification & multiplexed amplicon sequencing with ONT</td><td>epdl2</td><td>1779</td><td>ON863834</td></tr>
<tr><td>Gymnarchus niloticus</td><td>Unannotated genome assembly available (NCBI bioproject PRJNA423259)</td><td>epdl2</td><td>5280</td><td>ON863822</td></tr>
<tr><td>Ivindomyrus marchei</td><td>DNA extraction, PCR amplification & multiplexed amplicon sequencing with ONT</td><td>discarded[b]</td><td>-</td><td>-</td></tr>
<tr><td>Marcusenius moori</td><td>DNA extraction, PCR amplification & multiplexed amplicon sequencing with ONT</td><td>epdl2</td><td>1808</td><td>ON863836</td></tr>
<tr><td rowspan="3">Marcusenius ntemensis</td><td rowspan="3">DNA extraction, PCR amplification & multiplexed amplicon sequencing with ONT</td><td>epdl2.1</td><td>1675</td><td>ON863842</td></tr>
<tr><td>epdl2.3</td><td>1823</td><td>ON863841</td></tr>
<tr><td>epdl2.4</td><td>1806</td><td>ON863840</td></tr>
<tr><td>Mormyrops zanclirostris</td><td>DNA extraction, PCR amplification & multiplexed amplicon sequencing with ONT</td><td>epdl2</td><td>1869</td><td>ON863859</td></tr>
<tr><td>Mormyrus rume</td><td>DNA extraction, PCR amplification & multiplexed amplicon sequencing with ONT</td><td>epdl2</td><td>1816</td><td>ON863833</td></tr>
<tr><td rowspan="3">Paramormyrops curvifrons</td><td rowspan="3">DNA extraction, PCR amplification & multiplexed amplicon sequencing with ONT</td><td>epdl2.1</td><td>1676</td><td>ON863829</td></tr>
<tr><td>epdl2.3</td><td>1823</td><td>ON863831</td></tr>
<tr><td>epdl2.4</td><td>1819</td><td>ON863830</td></tr>
<tr><td rowspan="3">Paramormyrops hopkinsi</td><td rowspan="3">DNA extraction, PCR amplification & multiplexed amplicon sequencing with ONT</td><td>epdl2.2</td><td>1824</td><td>ON863857</td></tr>
<tr><td>epdl2.3</td><td>1793</td><td>ON863856</td></tr>
<tr><td>epdl2.4</td><td>1821</td><td>ON863858</td></tr>
<tr><td rowspan="3">Paramormyrops kingsleyae (APA)</td><td rowspan="3">paralog-specific PCR amplification & Sanger-sequencing</td><td>epdl2.1</td><td>1675</td><td>ON863825</td></tr>
<tr><td>epdl2.2</td><td>1814</td><td>ON863824</td></tr>
<tr><td>epdl2.3</td><td>1809</td><td>ON863823</td></tr>
<tr><td rowspan="2">Paramormyrops kingsleyae (BAM)</td><td rowspan="2">DNA extraction, PCR amplification & multiplexed amplicon sequencing with ONT</td><td>epdl2.1</td><td>1675</td><td>ON863849</td></tr>
<tr><td>epdl2.2</td><td>1813</td><td>ON863847</td></tr>
</table>

| | | | | |
|---|---|---|---|---|
| | | *epdl2.3* | 1810 | ON863848 |
| *Paramormyrops* sp. MAG (Type I) | DNA extraction, PCR amplification & multiplexed amplicon sequencing with ONT | *epdl2.2* | 1823 | ON863850 |
| | | *epdl2.3* | 1825 | ON863851 |
| | | *epdl2.4* | 1815 | ON863852 |
| *Paramormyrops* sp. MAG (Type II) | DNA extraction, PCR amplification & multiplexed amplicon sequencing with ONT | *epdl2.2* | 1825 | ON863826 |
| | | *epdl2.4* | 1817 | ON863827 |
| *Paramormyrops* sp. NGO | DNA extraction, PCR amplification & multiplexed amplicon sequencing with ONT | *epdl2.1* | 1676 | ON863846 |
| | | *epdl2.3* | 1815 | ON863845 |
| *Paramormyrops* sp. OFF | DNA extraction, PCR amplification & multiplexed amplicon sequencing with ONT | *epdl2.3* | 1823 | ON863844 |
| *Paramormyrops* sp. SN2 | DNA extraction, PCR amplification & multiplexed amplicon sequencing with ONT | discarded[c] | - | - |
| *Paramormyrops* sp. SN3 | DNA extraction, PCR amplification & multiplexed amplicon sequencing with ONT | *epdl2.3* | 1824 | ON863843 |
| *Paramormyrops* sp. SN9 | DNA extraction, PCR amplification & multiplexed amplicon sequencing with ONT | discarded[c] | - | - |
| *Paramormyrops* sp. SZA | DNA extraction, PCR amplification & multiplexed amplicon sequencing with ONT | *epdl2* | 1820 | ON863828 |
| *Paramormyrops* sp. TEN | DNA extraction, PCR amplification & multiplexed amplicon sequencing with ONT | *epdl2.2* | 1819 | ON863854 |
| | | *epdl2.3* | 1805 | ON863853 |
| | | *epdl2.4* | 1817 | ON863855 |
| *Paramormyrops* sp. TEU | DNA extraction, PCR amplification & multiplexed amplicon sequencing with ONT | discarded[c] | - | - |
| *Petrocephalus simus* | DNA extraction, PCR amplification & multiplexed amplicon sequencing with ONT | *epdl2* | 1792 | ON863832 |
| *Pollimyrus adspersus* | DNA extraction, PCR amplification & multiplexed amplicon sequencing with ONT | *epdl2* | 1551 | ON863838 |
| *Scleropages formosus* | genome assembly available (Ensembl gene ENSSFOG00015021850) | *epdl2* | 2918 | - |
| *Stomatorhinus ivindoensis* | DNA extraction, PCR amplification & multiplexed amplicon sequencing with ONT | *epdl2* | 1808 | ON863839 |

[a]From start to stop codon. bp = base pairs
[b]This sample was likely an incorrectly identified *P. kingsleyae*
[c]No *epdl2* genes were successfully amplified and sequenced

Table S3. Additional information on the live specimens used, including the NCBI SRA identifiers of the sequencing reads obtained.
Note: this table is released as a standalone .xlsx file.

Table S4. Primers used to amplify and sequence *epdl2* genes across Mormyridae

| primer | sequence (5'-3') | target gene | target clade | PCR annealing temperature ($^{o}$C) |
|---|---|---|---|---|
| epdl2.1_1F | CAGCCAGTGCGTCTACCATTTGC | *epdl2.1* | *P. kingsleyae* | 71 |
| epdl2.1_1R | AGGAATGAAACGAACAAAAGTTCAGGCAAGT | | | |
| epdl2.2_2F | GGTGAACTGCAGGTCTAGTTTG | *epdl2.2* | *P. kingsleyae* | 65 |
| epdl2.2_2R | ACAGAAGTTCAGGCAACTTTAACTTC | | | |
| epdl2.3_2F | AACCTACAAGGGACTTTGCTAACCC | *epdl2.3* | *P. kingsleyae* | 64 |
| epdl2.3_1R | GCCATGGACTACTTCTACAGCGCAG | | | |
| epdl2_Sanger_1 | TTCCTATCTGCCCTGGTA | *epdl2.1, epdl2.2, epdl2.3* | *P. kingsleyae* | NA |
| epdl2_Sanger_2 | TATCGCTGGGATTTCGAG | *epdl2.1, epdl2.2, epdl2.3* | *P. kingsleyae* | NA |
| epdl2_Sanger_3 | CTGTTATCCTAGGGATGAGG | *epdl2.1, epdl2.2, epdl2.3* | *P. kingsleyae* | NA |
| epdl2_Sanger_4 | CTGTCCAGGTTCTAATGC | *epdl2.1, epdl2.2, epdl2.3* | *P. kingsleyae* | NA |
| epdl2_Sanger_5 | TTCCAGACAAGCTCACTG | *epdl2.1, epdl2.2* | *P. kingsleyae* | NA |
| epdl2_0408_F01 | AGCARCRACACATTTTTGK | all *epdl2* genes | *Ivindomyrus, Marcusenius ntemensis, Mormyrus, Paramormyrops* | 60[a] |
| epdl2_0408_R01 | AGGGTTTSGAGTCAGGR | | | |
| epdl2_Pol+Sto_F01 | GTTGTTTTCAAAGTCGTCC | all *epdl2* genes | *Pollimyrus, Stomatorhinus* | 60 |
| epdl2_Pol+Sto_R01 | TGAACCTTGGATTCACAC | | | |
| epdl2_Bre+Hyp_F01 | TTACTTTCAGTGCTGTATC | all *epdl2* genes | *Brevimyrus* | 53 |
| epdl2_Bre+Hyp_R01 | CAGAGAATGCAGATAATTCAC | | | |
| epdl2_Mar+Cam_F01 | GTTGTTTTCAAAGTCGTCC | all *epdl2* genes | *Campylomormyrus, Gnathonemus, Marcusenius* | 52, 53, 55[b] |
| epdl2_Mar+Cam_R01 | TCTCTCTCCCTCTGATATA | | | |
| epdl2_Morps_F01 | CGAATCCTTAAATCCCAATC | all *epdl2* genes | *Mormyrops* | 55 |
| epdl2_Morps_R0 | CTGTAACGATTCACATGA | | | |

| | | | | |
|---|---|---|---|---|
| 1 | C | | | |
| epdl2_Pet_F01 | AGGGACAAYTTAGTCAG GA | all *epdl2* genes | *Petrocephalus* | 60 |
| epdl2_Pet_R01 | TTGAGTCAGAGRACACA GT | | | |

NA: primer not used in PCR reactions
[a]Touch-up annealing temperatures for: *Paramormyrops curvifrons*, *Paramormyrops* sp. OFF, *Paramormyrops* sp. SN2, *Paramormyrops* sp. SN9, *Paramormyrops* sp. TEU. Touch-up conditions: one set was comprised of 7 cycles from 57 to 60°C in 0.5°C increments. Each PCR consisted of 4 sets followed by 4 additional cycles at 60°C.
[b]*Campylomormyrus*: 52°C, *Gnathonemus*: 55°C, *Marcusenius*: 53 °C

Table S5. PCR reagents and final concentrations used to amplify each *epdl2* gene in *P. kingsleyae*

| Reagent | Final concentration |
|---|---|
| 5x Q5 reaction buffer | 1x |
| dNTPs | 200 µM each dNTP |
| Forward and Reverse Primers | 1.0 µM each |
| Template DNA | <1000 ng |
| Q5 polymerase | 0.02 U/µl |
| 5x GC enhancer for Q5 | 0.5x |

Table S6. PCR conditions used to amplify each *epdl2* gene in *P. kingsleyae*

| Step | temperature (°C) | time |
|---|---|---|
| Preheat lid + block | 98 | - |
| Initial denaturation | 98 | 30 s |
| 30 cycles of: | | |
| denaturation | 98 | 10 s |
| annealing | primer-dependent (Table S4) | 25 s |
| extension | 72 | 2 min |
| Final extension | 72 | 2 min |

Table S7. Mormyrid species and their NBCI bioproject codes that guided *epdl2* primer design

| species | BioProject |
|---|---|
| *Boulengeromyrus knoepffleri* | PRJNA526756 |
| *Brevimyrus niger* | PRJNA526749 |
| *Genyomyrus donnyi* | PRJNA529465 |
| *Gnathonemus echidnorhynchus* | PRJNA529468 |
| *Hyperopisus bebe* | PRJNA529477 |
| *Isicthys henryi* | PRJNA529470 |
| *Ivindomyrus marchei* | PRJNA529476 |
| *Marcusenius schilthuisiae* | PRJNA529469 |
| *Mormyrops attenuatus* | PRJNA530793 |
| *Mormyrops boulengeri* | PRJNA530782 |
| *Mormyrops zanclirostris* | PRJNA530797 |
| *Mormyrus hasselquistii* | PRJNA542939 |
| *Mormyrus iriodes* | PRJNA542943 |
| *Mormyrus proboscirostris* | PRJNA530791 |
| *Myomyrus macrops* | PRJNA423275 |
| *Myomyrus pharao* | PRJNA547756 |
| *Paramormyrops hopkinsi* | PRJNA547741 |
| *Paramormyrops* sp. MAG | PRJNA547743 |
| *Petrocephalus microphthalmus* | PRJNA423286 |
| *Petrocephalus schoutedeni* | PRJNA547742 |
| *Petrocephalus sullivani* | PRJNA427158 |
| *Petrocephalus zakoni* | PRJNA547751 |
| *Pollimyrus isidori* | PRJNA547785 |
| *Pollimyrus plagiostoma* | PRJNA547754 |
| *Stomatorhinus walkeri* | PRJNA547748 |

Table S8. PCR reagents and final concentrations used to amplify all *epdl2* genes across Mormyridae

| Reagent | Final concentration |
|---|---|
| 5x Q5 reaction buffer | 1x |
| dNTPs | 200 µM each dNTP |
| Forward and Reverse Primers | 0.9 µM each[a] or 0.5 µM each[b] |
| Template DNA | <1000 ng |
| Q5 polymerase | 0.02 U/µl |
| 5x GC enhancer for Q5 | 0.1x |

[a]epdl2_0408 primers with *Ivindomyrus marchei*, *Marcusenius ntemensis*, and all *Paramormyrops* spp
[b]epdl2_0408 primers with *Mormyrus rume*, and all other primers

Table S9. PCR conditions used to amplify all *epdl2* genes across Mormyridae

| Step | temperature (°C) | time |
|---|---|---|
| Preheat lid + block | 98 | - |
| Initial denaturation | 98 | 30 s |
| 30[a] cycles of: | | |
| denaturation | 98 | 10 s |
| annealing | primer- and species-dependent (Table S4) | 20 s[b] |
| extension | 72 | 75 s[c] |
| Final extension | 72 | 2 min |

[a]32 cycles for: *Brevimyrus niger*, *Campylomormyrus* sp*, Ivindomyrus marchei*, *Marcusenius ntemensis*, *Paramormyrops curvifrons*, *Paramormyrops* sp. OFF, *Paramormyrops* sp. SN2, *Paramormyrops* sp. SN9, *Paramormyrops* sp. TEU

[b]25 seconds for: *Brevimyrus niger, Campylomormyrus* sp*, Ivindomyrus marchei*, *Marcusenius ntemensis*

[c]90 seconds for: *Brevimyrus niger, Campylomormyrus* sp

Table S10. Sites along Epdl2 that have experienced positive selection in the osteoglossiform *epdl2* gene tree and have evolved at increased ω rates in mormyrid lineages with vs without *epdl2* duplications. Branches supported by each EBF value are marked by rectangles in Fig. S3 (EBF values for a given site are arranged left to right to match their corresponding branches from top to bottom)

| Site | Estimated number of branches under positive selection | Empirical Bayes Factor (EBF) |
|---|---|---|
| 28 | 1 | $1.1 \times 10^{4}$ |
| 105 | 1 | $2.6 \times 10^{11}$ |
| 106 | 0 | - |
| 125 | 1 | $5.5 \times 10^{4}$ |
| 126 | 2 | 1449, $1.0 \times 10^{26}$ |
| 127 | 2 | 2200, $3.2 \times 10^{10}$ |
| 129 | 2 | 543, $1.0 \times 10^{26}$ |
| 150 | 2 | 2165, 315 |
| 153 | 1 | $8.8 \times 10^{12}$ |
| 172 | 4 | $2.4 \times 10^{4}$, $2.4 \times 10^{4}$, $1.0 \times 10^{26}$, 1043 |

Table S11. Summary of the amino acid substitutions observed in the Epdl2 paralogs at the ten sites under positive selection and increased ω rates in mormyrid lineages with vs without *epdl2* duplications

| site | ancestral amino acid | derived amino acid residues observed in each paralog (total) | | | |
|---|---|---|---|---|---|
| | | *epdl2.1* (5) | *epdl2.4* (6) | *epdl2.2* (6) | *epdl2.3* (10) |
| 28 | S | - | - | P (4) | P (10) |
| 105 | F | - | - | S (1) | - |
| 106 | P | R (5) | L (6) | R (6) | R (10) |
| 125 | S | - | N (1) | N (6) | N (10) |
| 126 | S | - | - | - | L (8) |
| 127 | A | - | - | D (6) | - |
| 129 | G | - | S (1) | S (2) | S (2) |
| 150 | Q | L (5) | L (4), K (2) | K (6) | K (10) |
| 153 | F | - | - | C (1) | - |
| 172 | L | - | - | R (1) | W (7), R (2) |

### Figures

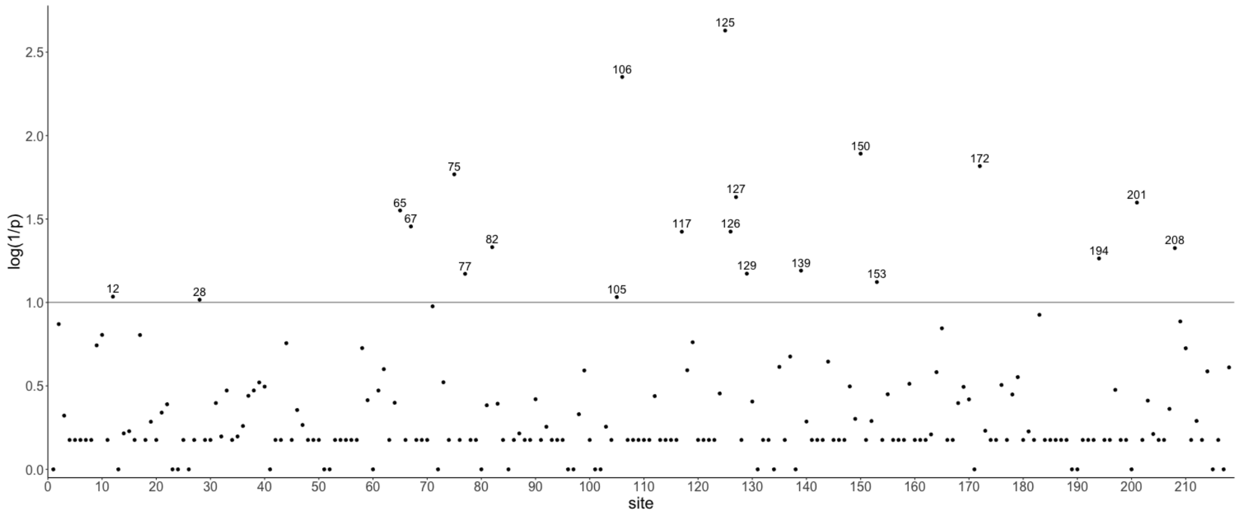


Fig. S1. Sites along Epdl2 that have experienced positive selection (MEME, $p < 0.1$) in the osteoglossiform taxa studied. p values have been transformed so that higher values on the y axis represent lower p values. Horizontal line marks the significance threshold and significant sites are labeled.

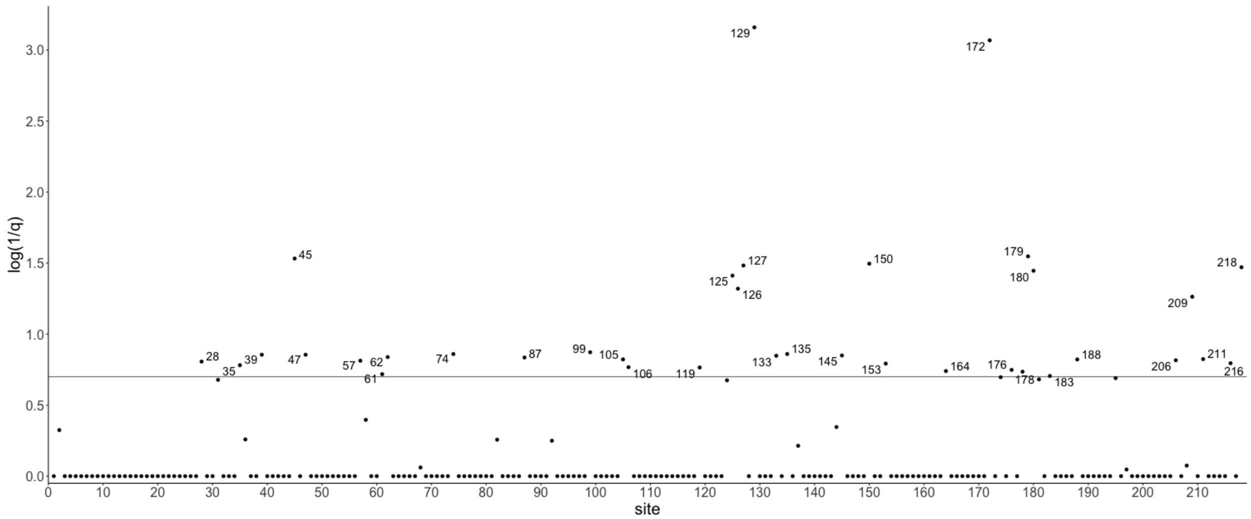


Fig. S2. Sites along Epdl2 with higher ω values in mormyrid lineages with vs without epdl2 duplications (Contrast-FEL, $q < 0.2$). q values have been transformed so that higher values on the y axis represent lower q values. Horizontal line marks the significance threshold and significant sites are labeled.

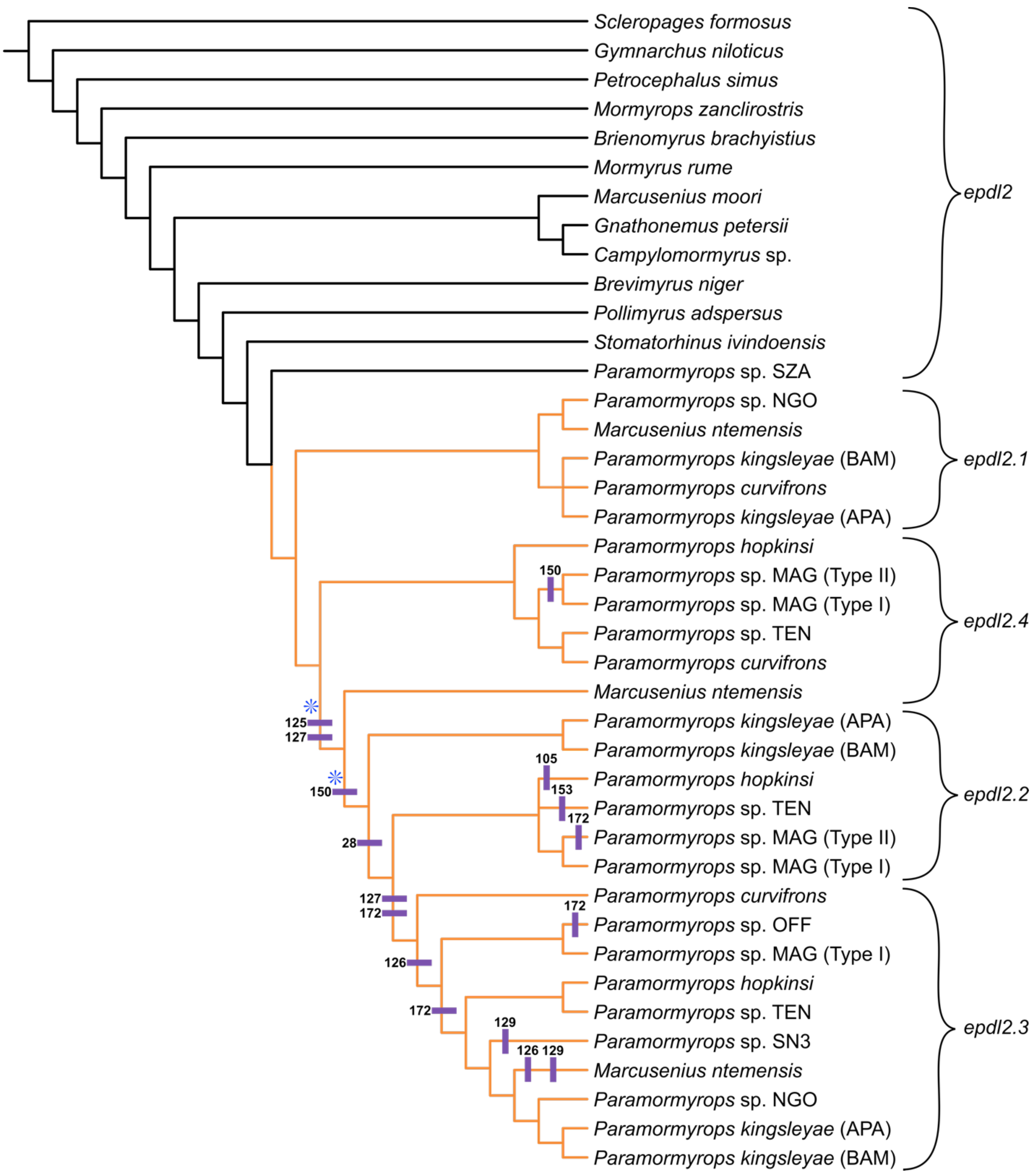


Fig. S3. Topology of the osteoglossiform *epdl2* gene tree, we highlight branches and sites where we detected signals of selection. Orange branches are recently duplicated *epdl2* paralogs, selection has intensified in these branches relative to the mormyrid lineages without *epdl2* duplications. Branches with blue asterisks experienced positive selection. Purple rectangles are labeled with sites along Epdl2 where selection has intensified in mormyrid lineages with *epdl2* duplications and have experienced positive selection. These rectangles are placed on the branches where exploratory evidence suggests they underwent positive selection. All rectangles map to the lineages with *epdl2* duplications.
